# Variation in protein coding genes identifies information flow as a contributor to animal complexity

**DOI:** 10.1101/679456

**Authors:** Jack Dean, Daniela Lopes Cardoso, Colin Sharpe

## Abstract

Across the metazoans there is a trend towards greater organismal complexity. How complexity is generated, however, is uncertain. Since *C.elegans* and humans have approximately the same number of genes, the explanation will depend on how genes are used, rather than their absolute number. Functional diversity is a measure that quantifies the isoforms, domains and paralogues of a gene. In this paper we determine functional diversity for each protein-coding gene in the human genome and its orthologues across eight commonly used model organisms. From this we derive the Compx list of genes that correlate positively with their increase in cell-type number. We then select genes common to the Compx list and the ExAC list, whose genes show minimal variation across many human genomes, and identify genes with common functions, notably in chromatin structure, RNA splicing, vesicular transport and ubiquitin-mediated protein degradation. Together these functions reveal that information flow within the cell relates to cell-type number, used as a measure of organismal complexity.

## Introduction

Some animals are more complex than others. For example, the nematode worm, *C. elegans*, widely used as a model organism, has many fewer cell types than vertebrate models, such as the mouse. Consequently, cell-type number has been widely applied as a quantitative measure of complexity^1 2^. Since the number of genes in the genome does not correlate with complexity^3^, it is more likely that patterns of gene expression and the nature of the encoded proteins are important determinants.

Patterns of gene expression are determined by cis-acting regulatory elements^4 5^. Since these frequently act independently, the loss or gain of an element will have little effect on the expression pattern of the gene mediated by other elements. Cis-acting regulatory elements can therefore confer the capacity for adaptive evolution as documented by the different spatial patterns of the Pitx1 expression in sticklebacks that cause a variation in body armour, dependent on the environment ^6^ and their role in the loss of powered flight in some birds^7^. Although the Encode project ^8^ has mapped many cis-acting regulatory elements within the human genome, comprehensive data is lacking for most other species. Instead, in this paper we focus on a set of factors that influence protein function, to analyse how changes to these factors in orthologues relate to the complexity of multicellular animals.

Several of these factors have been shown to correlate positively with the increase in cell-type number (CTN) across the metazoan phylogeny. For example, *C. elegans* produces on average just over 1 transcript type per gene (protein coding and non-coding), whilst in humans this rises to around 5 (based on Ensembl^9^ assembly statistics) through the wider application of alternative splicing and the use of multiple transcript start sites. The resulting transcript diversity has been positively correlated with animal complexity^2^. In addition, the complement of protein domains and motifs within a protein across a range of species has been shown to correlate with complexity, again as measured by CTN^1 10^. Although the overall number of genes is unlikely to contribute, variation in the number of related genes, the paralogues within a species, could also be important^11^. When a gene is duplicated the resulting daughter genes have the opportunity either to take on new activities (neofunctionalization)^12^ or to divide the activities between the two new genes (subfunctionalisation)^13^.

Combining data on the number of paralogs, transcript isoforms and protein domains associated with a gene defines a value termed the functional diversity (D_F_) of the gene^14^. Correlating D_F_ with cell-type number across a range of metazoans for over 2000 genes associated with transcription identified genes involved in chromatin dynamics as candidate contributors to animal complexity^14^.

This paper determines the functional diversity for each annotated gene in the human genome and then across an additional eight species commonly used as model organisms for biomedical research (Supplementary Data 1). This approach extends previous analysis to the complete human gene list and its orthologues generating data for more than 120 000 genes to identify a list of genes, the Compx list, whose function may underpin complexity. We then identify a prominent overlap between the Compx list and the ExAC list of human genes that show reduced levels of variation within the human population^15^. It is thought that the ExAC genes are under purifying selection because they encode proteins that play important roles within the cell. Analysis of the overlapping genes found in both lists indicates that many encode proteins that determine information flow within the organism, relating increases in cell-type number to the ability to process information. This defines a wide, systems-based context in which to evaluate the contribution of specific genes and cellular processes to animal complexity.

## Methods

### Data-mining and calculation of functional diversity

For each human gene, information on the number of paralogues was obtained from the Ensembl database^9^. Similarly, the number of protein coding transcripts and the number of protein domains encoded by each gene, as defined by the Ensembl Prosite Profiles^16^, was extracted from the Ensembl database using Biomart and adapted protocols (https://github.com/JackDean1/OrganismComplexity). The pipeline was then used to extract the same information for orthologues of the human genes across eight additional species from the nematode worm, *C. elegans* to the Macaque, *Macaca mulatta*. Since not all human genes have orthologues across all the species, the dataset covers 19 908 identifiable human genes (97.5% of the Ensembl human gene list, unmapped genes were excluded) and 127 355 genes in total. Details of the genomes analysed are provided in Supplementary Data 1.

A value for the functional diversity (D_F_) of each gene was then calculated. The protocol is essentially as described^14^ and is calculated from:

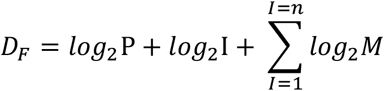

Where P is the number of paralogues, I the number of isoforms and M the number of protein motifs and domains associated with the gene. The complete dataset and calculation of D_F_ is in Supplementary Data 2.

### Correlation analysis

Ortho-sets, which are named after the human gene, were selected that contained the human gene and either eight, seven or six orthologues allowing for the absence, or lack of annotation, of some genes in some species. The D_F_ values for each gene within the ortho-set were correlated with cell type number (CTN)^1 2 17^, taken as a measure of the complexity of an organism, using a Pearson’s correlation (r) that takes account of the degrees of freedom defined by the number of orthologues in each ortho-set. A similar approach was taken to identify ortho-sets that showed only minimal change in the value of D_F_ for each gene of the ortho-set. Lists of ortho-sets from the ‘complexity’ analysis and the ‘minimal change’ analysis were compiled with increasing statistical probability in a two-tailed t-test for the Pearson’s correlation and the overlap between the lists determined. At p<0.05 there was a 10% overlap in the ortho-sets between the two lists, but this reduced to a 0.2% overlap at p<0.01. Consequently, the complexity ortho-set list at p<0.01, called the Compx list, was used in all analyses.

### GO-term and STRING analysis of the gene lists

The Compx list of 571 genes (Supplementary Data 3) was subject to a Gene Ontology (GO)-term over-representation analysis using Amigo2 v2.5.12^18^ and the Panther Classification system v14^19 20^ to identify terms enriched by more than 2.5 fold and with an adjusted probability of less than 0.05 (Fisher’s Exact test with Bonferroni correction for multiple testing). Redundancy between terms within and across the three major GO-terms was identified on a Circos plot (gene lists in Supplementary Data 4). Interactions between the 571 Compx genes were identified using the STRING application^21^.

### Validation of the Compx – ExAC list overlap

A bootstrap approach was taken to determine whether the overlap between the Compx and ExAC gene lists was greater than might be expected at random. Using simply the numbers of genes in each list (571 for Compx, 3230 for ExAC and 19 908 for the human genome) the overlap between 571 random numbers between 1 and 19 908 and 3 230 random numbers between 1 and 19 908 was determined across 1000 iterations and the mean and standard deviation calculated. The process was then repeated replacing 571 with 362, the number of genes in the ‘no change’ analysis. These calculated, expected values were then related to the actual overlaps between the gene lists.

Keywords associated with the function of each of the 159 Compx-ExAC overlap genes were extracted from GeneCards^22^ and validated by reference to published information. The gene list was then compiled by major function (Supplementary Data 7).

## Results

### Creating a database of functional diversity for human protein coding genes and their orthologues in eight metazoan species

This analysis is based on the premise that changes to the biological activity of proteins will be one of a number of factors that contributes to organismal complexity^1 11 14 23^. When comparing orthologous genes, the range of biological activity will depend on three factors; the number of different protein-coding transcripts that can be generated from each gene, the types and numbers of each domain that are encoded in the protein and the number of paralogues present in the genome^14^. Values for each of these factors were extracted from Ensembl, using Biomart, to produce a database for 19 908 human protein coding genes and 127 355 protein coding genes across the nine species considered (Supplementary Data 2) representing a comprehensive coverage of genes from many of the model organisms used in biomedical research. The values were then used to calculate the functional diversity (D_F_) of each gene in each species, providing a metric for the capacity of that gene to provide the diverse biological activities that determine complexity. As described in the Methods section, correlating the change in value of D_F_ with the change in value of the cell type number, taken as a measure of organismal complexity, identified a list of 571 ortho-sets that forms the Compx list

### GO-term over-representation analysis identifies a limited number of activities

Whilst it is possible that any one of the 571 ortho-sets may contribute to animal complexity, to identify Compx those with common features, we compared the Compx and human gene lists to identify GO-terms over-represented in the Compx list. The Panther Classification system^19 20^ was used to identify GO-terms that are over-represented more than 2.5 fold and with an adjusted p-value of less than 0.05. After the resolution of redundant, nested terms, seven GO-terms met these criteria (Fig. 1 and Supplementary Data 4). These included two associated with the processing of alcohol, two involving mRNA processing, including the spliceosome complex, and one each for transcription-coupled nucleotide excision repair, Golgi vesicle transport and the actin cytoskeleton. To further analyse the over-represented genes, the degeneracy between the ortho-sets both within and across the three primary GO-terms was determined (Fig. 1B, C) to identify a definitive list of 109 ortho-sets (Supplementary Data 4). Combined with a Panther over-representation test for the reactome (Fig. 1D) this identified four main activities: ethanol oxidation, the regulation of splicing, DNA repair and membrane trafficking, specifically of ER-Golgi vesicles.

**Figure 1.**
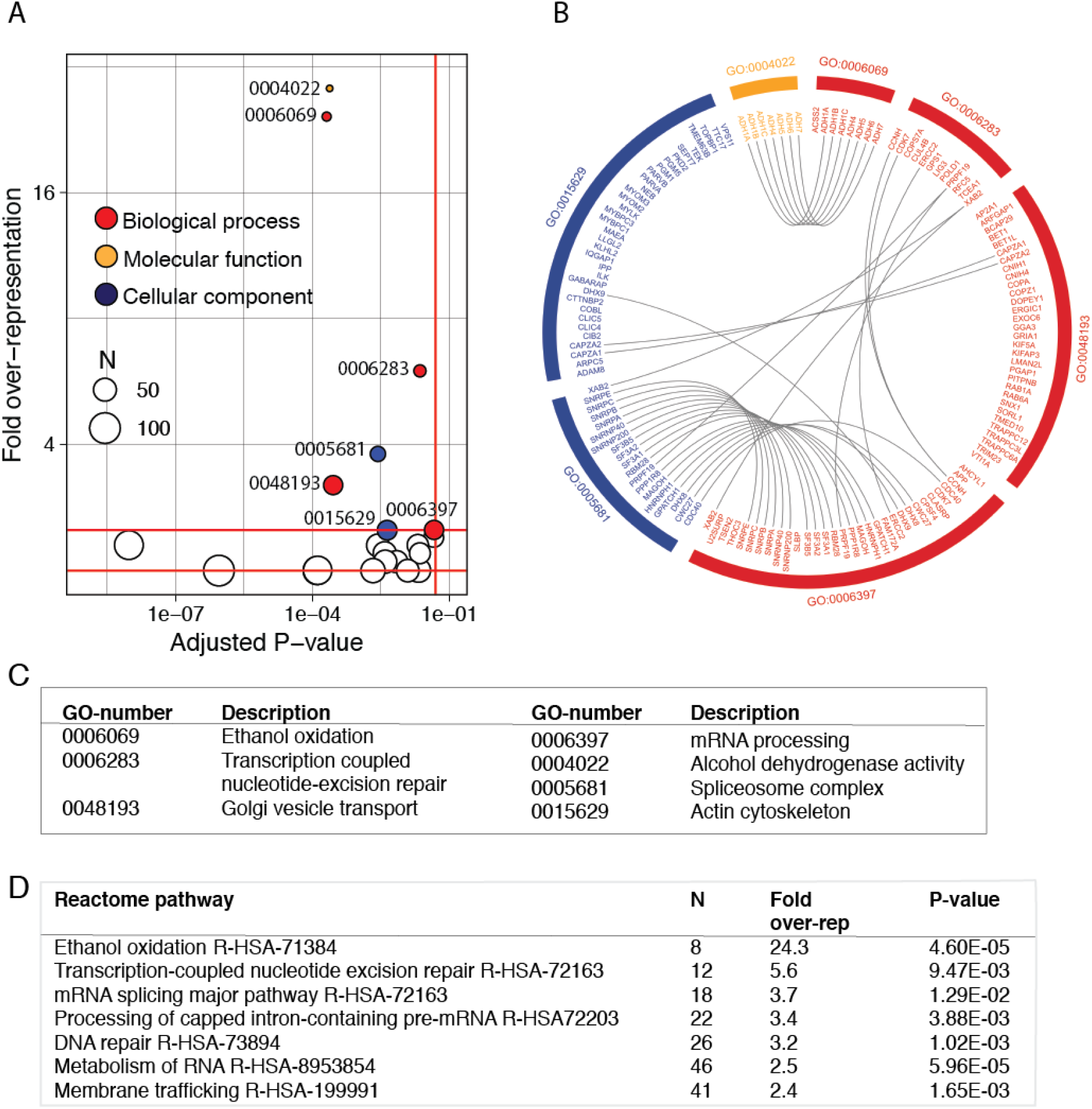
Over-representation assay for the 571 Compx list. A. Combined graph for the three primary GO-term fields. 2-fold and 2.5-fold over-representation levels are shown and terms that meet the required criteria are coloured according to the primary GO-term. Terms with >2<2.5-fold over-representation are shown as open circles and are listed in Supplementary Data 4. B. Circos plot showing redundant ortho-sets between the GO terms. Complete lists of ortho-sets are in Supplementary Data 4. C. Identification of the major GO-terms. D. Identification of the major terms within the Panther Reactome for over-represented genes within the Compx list.

### Identifying interacting subsets using STRING analysis

Identifying interactions between proteins encoded by the Compx list will point to additional functions to which these genes contribute. The bioinformatic STRING program^21^, set to identify high confidence, experimentally determined interactions identified 7 activities across 4 networks. (Fig. 2).

**Figure 2:**
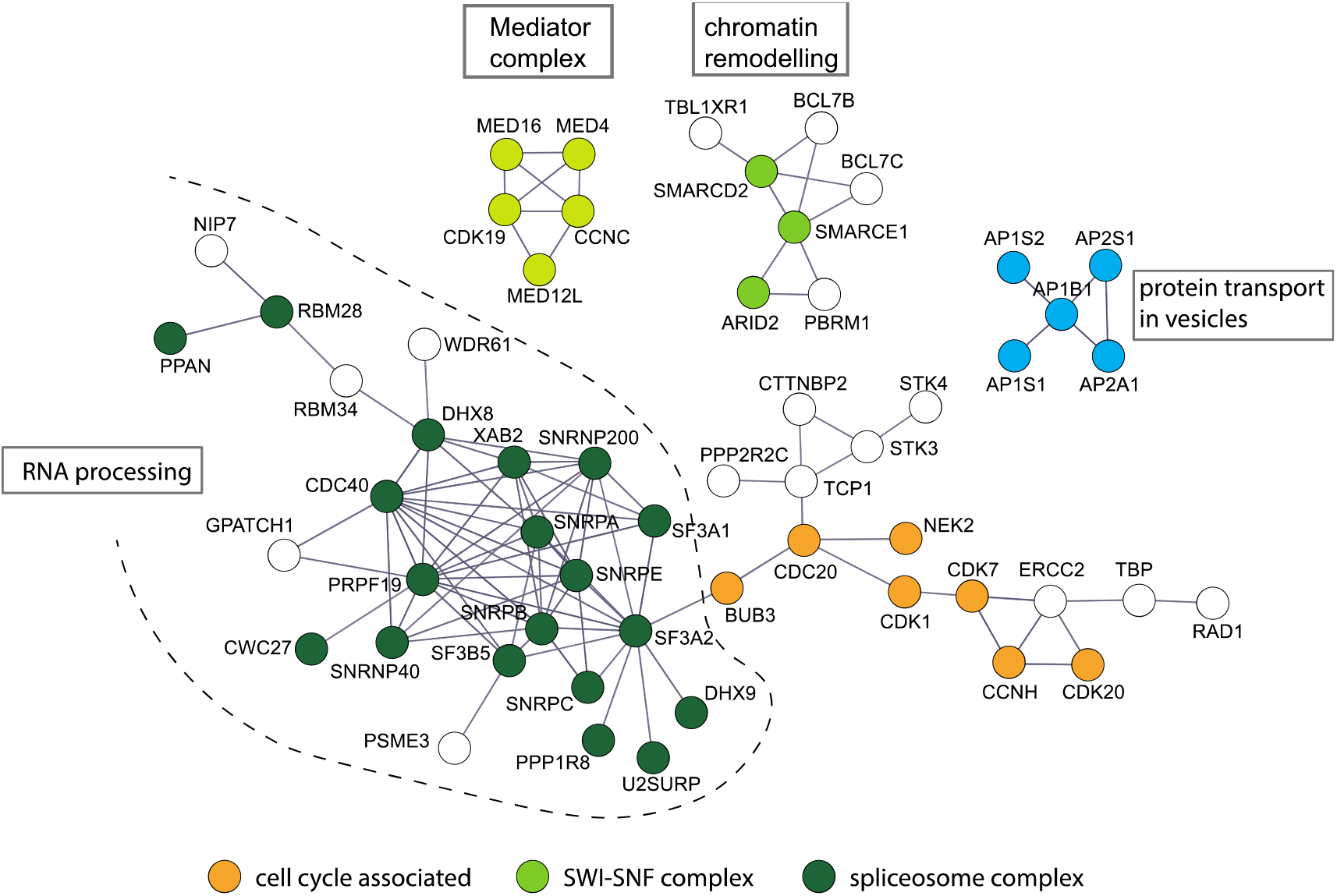
Interaction analysis of the Compx ortho-set. Only experimentally verified (confidence level 0.7) interactions and those networks containing at least one node with three connections are shown. Networks have been coloured by function. The distance between genes and the orientation is random. The analysis identified 7 major functions across 4 networks.

Conserved activities between the two approaches include RNA splicing (the regulation of splicing and snRNP complex) and protein-vesicle transport (associated mainly with the ER-Golgi). In addition, the interactions analysis identified members of the SWI/SNF chromatin remodelling complex, proteins that impose repressive chromatin conformations, the Mediator complex and proteins that contribute to the cell cycle.

### Filtering the Compx list to identify genes with a critical function

The ExAC genes are identified as the human genes that, following sequencing of over 60 000 genomes, show less than expected variation^15^. The suggestion is that the function of genes on the ExAC list cannot be maintained if the gene is subject to the standard rate of mutational change, indicating that their products may play critical roles in the cell^15^. We then hypothesise that genes from the ExAC list may be over-represented in the Compx list. This would then provide a filter to identify genes that are both vital to the cell and contribute to animal complexity.

A direct comparison shows that 159 of the ExAC list of 3230 genes are represented in the 571 gene Compx list (Supplementary Data 5). A bootstrap method of 1000 sampling events shows that the expected overlap between two groups the size of the Compx and ExAC lists, selected at random from a pool the size of the human gene list, is 92. The observed figure of 159 genes is almost 8 standard deviations greater than the expected mean for the random selection (Fig. 3A).

**Figure 3:**
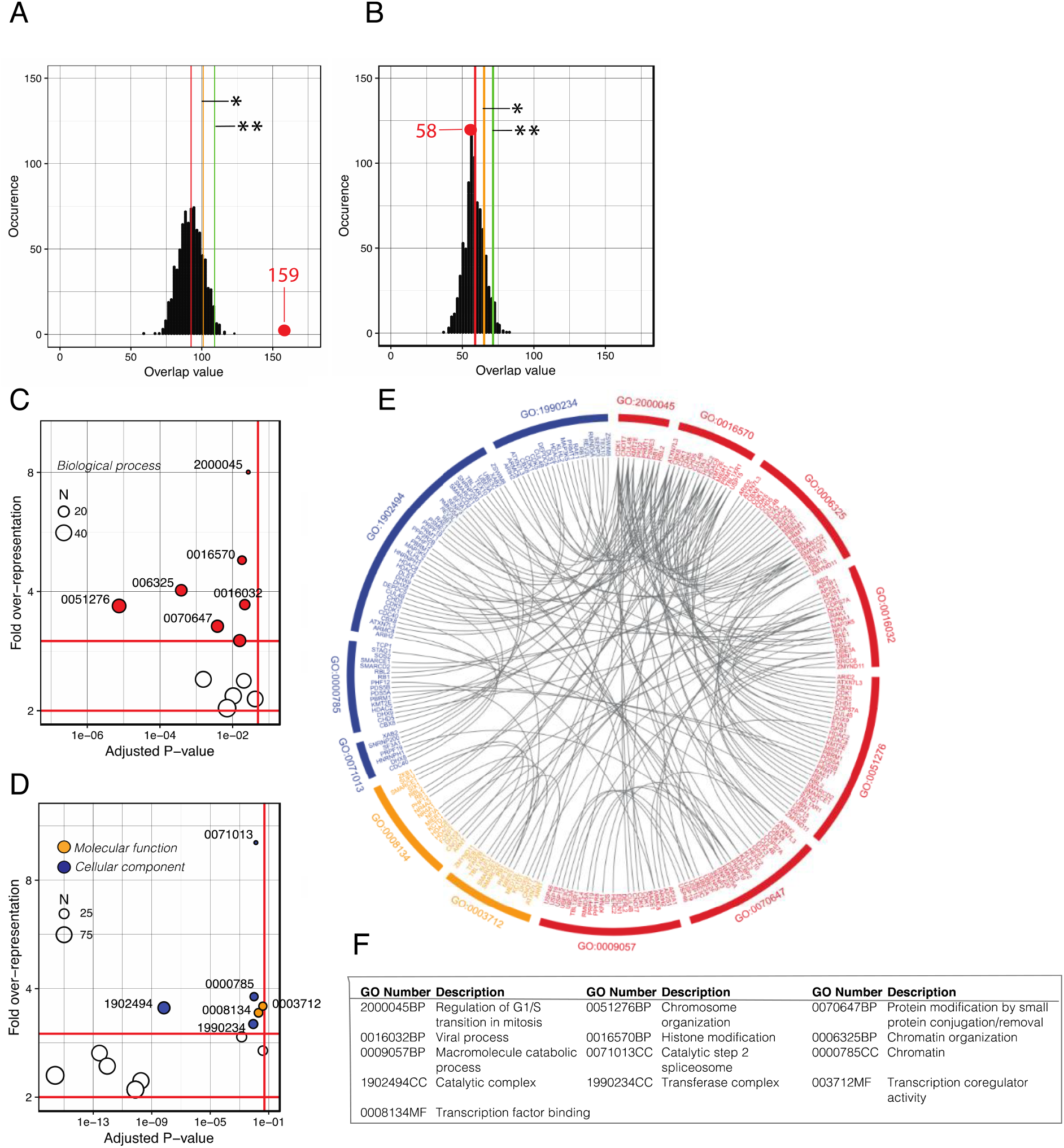
Demonstration of a prominent overlap between the 571 complexity genes set and ExAC list. A. The distribution of 1000 predicted overlap values between samples of size 571 corresponding to the complexity ortho-set list and 3230 corresponding to the ExAC gene list taken from a stock of 19908, corresponding to the number of human genes in the analysis. The actual overlap value 159 is marked by the red dot. B. The same analysis, repeated using a sample size of 362 corresponding to the minimal-change ortho-set. In each, the mean is marked by a red line, one standard deviation from the mean by the amber line (*) and two standard deviations by the green line (**). C. GO-term over-representation analysis for the complexity and ExAC overlapping gene list. For the analysis of the general GO-term, Biological Process, the red circles identify terms over-represented by >3x and with adjusted P<0.05. D. Combined analysis for the general GO-terms Cellular Component (blue circles) and Molecular Function (orange circles) (over-represented by >3x and adjusted P<0.05). The identity of the terms over-represented by >2<3x are listed in Supplementary Data 6. E. A Circos plot of the individual genes identifies a high degree of redundancy between the over-represented GO-terms and a unique list of 90 genes (Supplementary Data 6). F. Table of the over-represented GO-terms, the colours refer to the major GO-terms as shown in C and D where red denotes Biological Process, orange is Molecular Function and blue is Cellular Component.

In contrast, the same approach, replacing the Compx list with the 362 minimal-change ortho-set, described above, predicts an overlap with the ExAC list of 58 genes, which is identical to the observed figure of 58 (Fig. 3B). This approach validates the hypothesis of a relationship between genes of the Compx and ExAC lists and produces an overlap-list of 159 genes. Many of the genes resulting from this filter are likely to have both a crucial function in the cell and critically influence animal complexity.

### Analysis of the 159 genes that are common to the Compx and ExAC gene lists

The 159 overlapping genes were subject to GO-term over-representation analysis to highlight commonalities and the hierarchical terms rationalised using Revigo^24^. GO-terms were selected that met the criteria of a greater than 3-fold enrichment, when compared to the standard human genome gene list, and with a probability of less than 0.05 (Fig. 3C and D). This identified terms across the three major GO-term categories, notably associated with chromatin dynamics, transcription coregulation, RNA splicing and the cell cycle. In addition are the terms ‘protein modification by small protein conjugation or removal’ and ‘macromolecule catabolic process’ that refer extensively to protein ubiquitination and related processes including the breakdown of proteins in the proteasome^25^ and the elimination of misfolded proteins in the ER-Golgi system by ERAD^26 27^ (Fig. 4).

**Figure 4.**
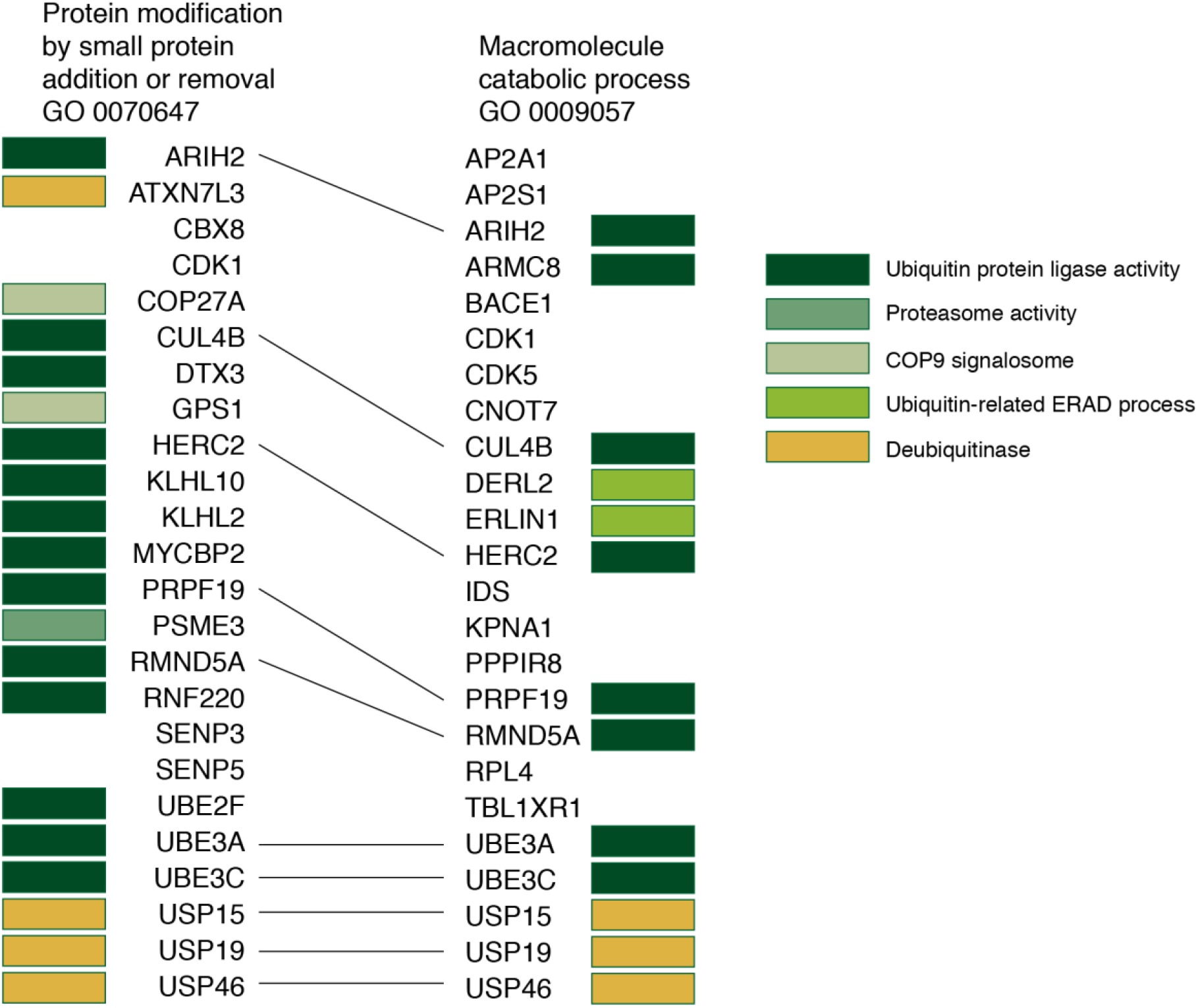
The overlap list contains ortho-sets associated with protein ubiquitination Isolating the overlap between GO:0070647 protein modification by small protein conjugation or removal and GO:0009057 macromolecule catabolic process, identifies genes involved in the tagging of proteins with ubiquitin and their recognition for breakdown in the proteasome and for the removal of misfolded proteins from the ER-Golgi by the ERAD process.

The extensive redundancy of genes within the GO-terms was plotted using Circos and identifies that 90 of the 159 ortho-sets (61%) are registered in the over-representation GO-term analysis (Supplementary Data 6).

The products of the 159 overlapping genes were analysed for protein interactions using STRING (Fig. 5).

**Fig 5.**
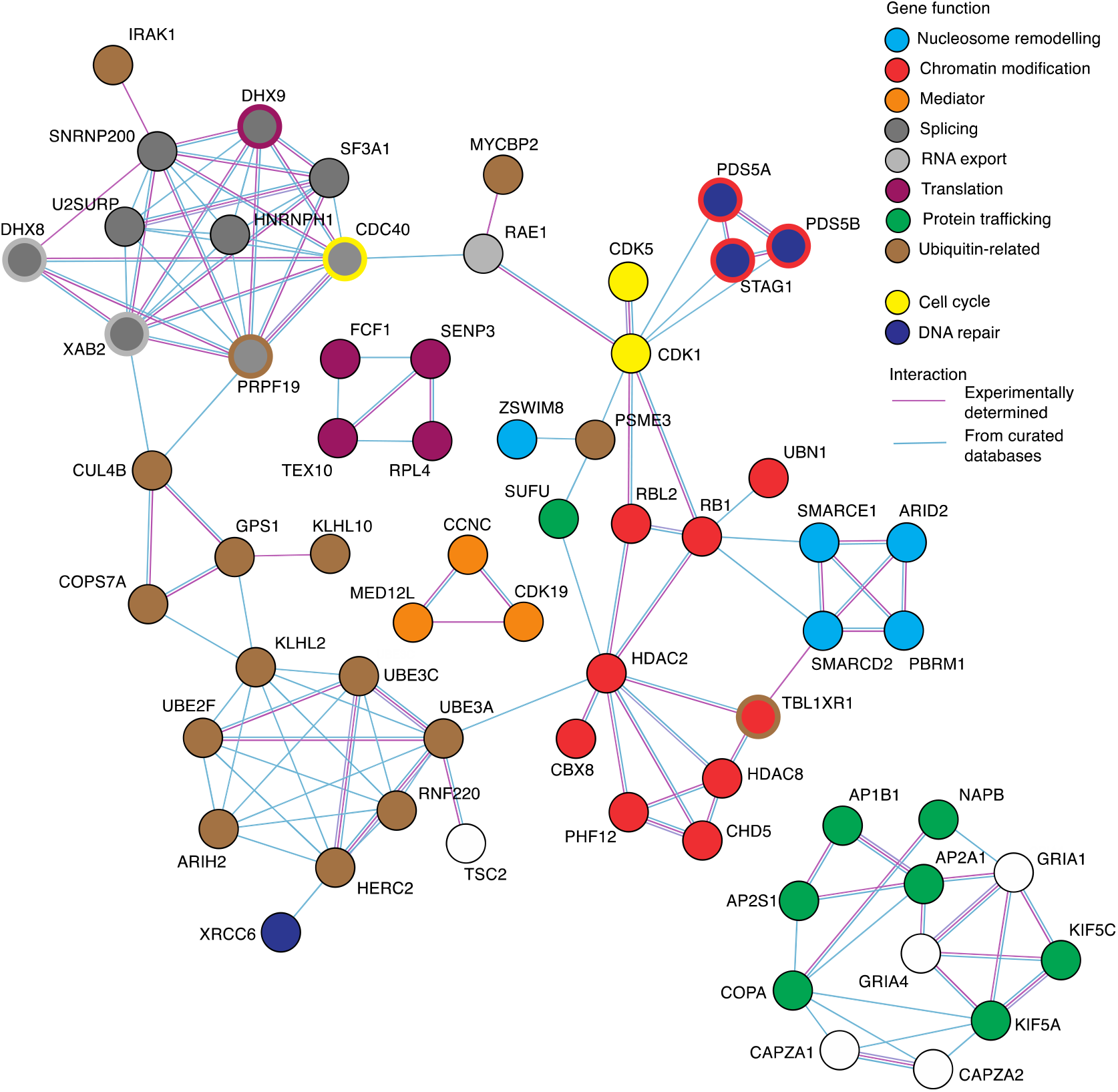
Interaction analysis of the Compx versus ExAC overlap gene list. Only networks with nodes that contained at least three connections are included. Nodes have been coloured to indicate function. The connectivity of the interacting nodes is more than expected for a random selection of genes with a p=8.0E-06.

STRING analysis identified a range of interacting groups that could be described in terms of the function of the genes, as described in GeneCards and identified eight major and two minor groupings (Fig. 5). Although there may seem to be little in common between the groupings, a closer consideration suggests these are all components that contribute to information flow in the cell, from access to the chromatin to the final removal of the protein. Information flow in the cell can be described as a sequence of seven events (Fig. 6 and see Discussion). In addition, the integrity of the information needs to be maintained by DNA repair mechanisms and information flow integrated with the cell cycle.

**Figure 6.**
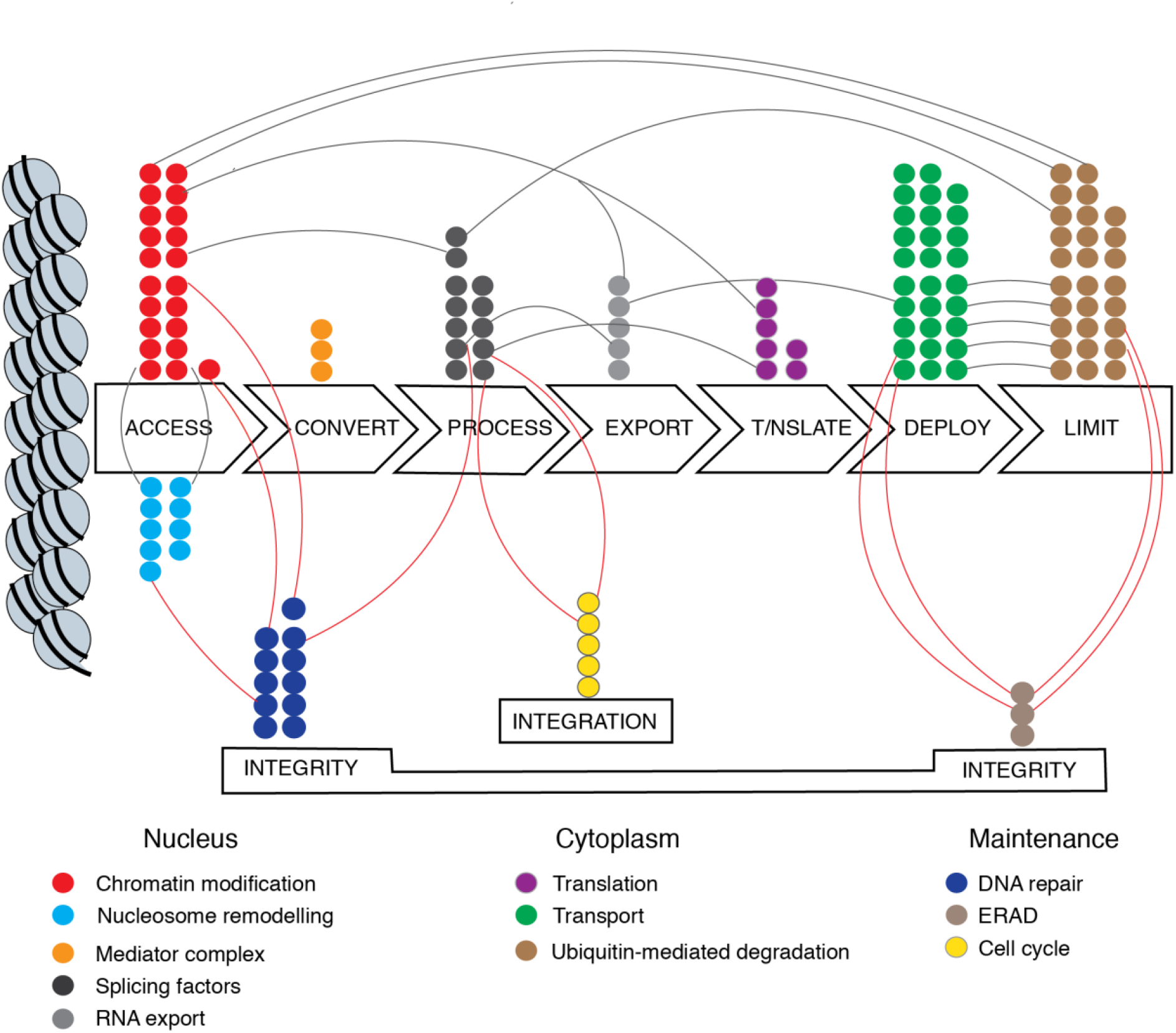
Mapping the Compx-ExAC overlap genes to the elements of information flow in the cell. Individual genes (coloured circles) are mapped to the seven elements of gene flow and to the two maintenance factors, integrity and integration. Justification for the allocation is given in Supplementary Data 7. The contribution of one gene product to more than one element is marked by a connecting arc.

To address further the possibility that complexity relates to information flow, the functions of the genes identified in the GO-term analysis that did not appear in the STRING analysis were similarly categorized by reference to GeneCards^22^ and published data. In total 81 of the 97 genes could be allocated to events relating to information flow. Of the remaining 61 genes, 25 could be similarly allocated, giving a total of 106 out of 159 genes (67%) that are held in common between the Compx and ExAC gene lists that may play a role in information flow in the cell (Supplementary data 7). These genes were mapped to the seven events (Fig. 6) and those genes that contribute to more than one event indicated. In addition to DNA repair, the integrity of information flow is maintained by the removal of mis-folded proteins in the ER-Golgi system through endoplasmic reticulum-associated degradation (ERAD)^26 27^.

These results indicate an important relationship between information flow and a cell-type number, a measure of complexity for multicellular organisms.

## Discussion

In this paper, we use a simple approach to data-mine three variables that define and quantify the functional diversity (D_F_) of proteins derived from each human gene and its orthologues from up to eight other species (termed an ortho-set). The change in D_F_ for each gene is correlated with the change in number of different cell types in the species, a proxy for organismal complexity, to identify 571 ortho-sets with a positive correlation.

This approach subscribes to a specific model in which incremental changes in functional diversity across an ortho-set relate to an incremental change in complexity. Alternative models, in which, for example, major changes in functional diversity occur at a specific point in the phylogeny, are also likely and can be extracted from our data set (Supplementary Data 2). Whilst individual genes, acting in isolation, are likely to relate complexity in either model, GO-term over-representation and interaction analysis point towards processes and systems and it is these that will contribute to a broader understanding of the basis of organismal complexity.

This paper considers functional diversity, but this will be only one of several factors associated with animal complexity. Amongst others, an increasing diversity in cis-acting regulatory elements (CAREs) across a phylogeny will play a role^4 5 28^ and there may also be a contribution to complexity from changes to the complement of regulatory non-coding RNAs, including the miRNAs^29 30^ and lncRNAs that have been shown to modulate gene expression^31^.

We have not been able to include all the factors that affect protein functional diversity in this analysis. For example, short linear motifs (SLiMs)^23^ impact on the function of proteins such as NCoR2 ^11^, but these are not currently widely annotated. The functional capabilities of a protein may also be honed by a small number of changes within the sequence that affect neither protein domain number, nor the range of isoforms^32^. Lack of this data may mean that some ortho-sets that contribute to complexity have not been identified.

Other studies have successfully identified the contribution of individual components to complexity, such as protein domains and alternative splicing^1 2^. A previous paper used the approach taken here to assess genes broadly associated with transcription, as listed in the AnimalTFDB 2.0 database of 2087 human genes^14^. The identified ortho-sets that are primarily involved in the dynamic structure and function of chromatin, including nucleosome remodeling, the modulation of chromatin activity and the Mediator function. By extending the analysis to the whole genome we aimed to expand this selection to include events outside the nucleus. GO-term over-representation analysis (Fig. 1A) indicated that the oxidation of alcohol, mRNA processing, notably splicing, aspects of DNA repair and protein transport via vesicles should also be considered. For the oxidation of alcohol, the main component of functional diversity is the reiterative gene duplication followed by sub- and neo-functionalization that have produced a range of enzymes that metabolize ethanol and related alcohols^33 34^. Analysis of interactions amongst the Compx ortho-sets via STRING identified components of the spliceosome and of vesicular transport and in addition highlights chromatin remodeling and the Mediator complex, consistent with the results from the previous, more limited, survey^14^.

In this paper we apply an additional filter by identifying the overlap between the Compx and ExAC genes lists. The ExAC genes have less than expected variation within the human population, suggesting they play important functions that are sensitive to mutation^15^. This might seem to contradict the essence of the Compx list, which selects for increases in functional diversity, but the Compx list reflects changes between orthologues across a wide phylogeny, rather than variation within a single species. There is a greater than expected overlap between the Compx list and the ExAC list that is not seen when the ExAC list is compared to the ‘minimal-change’ list of genes selected for minimally changing values of functional diversity (Fig. 4). The hypothesis is that the filter will highlight those ortho-sets that both relate to complexity and are important to the function of the cell, allowing the identification of cellular processes that are critical for organismal complexity.

Within the overlap list, GO-term analysis identified sub-terms associated with chromatin dynamics, the spliceosome, protein modification by ubiquitination and the cell cycle. Interaction analysis similarly identified ortho-sets associated with mRNA splicing and chromatin dynamics, notably nucleosome remodeling, ubiquitin-mediated protein turnover and, in addition, ortho-sets that encode proteins involved in the distribution of proteins via vesicles within the ER-Golgi apparatus.

Is there a common theme? Each of these groupings can be directly related to the concept of information flow in the cell. We hypothesise that the contribution of protein functional diversity to complexity is primarily, though not exclusively, at the level of information flow in the cell. Information flow can be explained in terms of seven sequential elements (Fig. 6). First, nucleosome re-modelling and epigenetic marks regulate *access* to chromatin, flipping chromatin between states that either promote or repress gene expression^35 36 37 38 39^. The Mediator complex then promotes interactions between distant enhancer elements and the genes to be expressed^40 41^ which in combination with transcription factors and RNA polymerases will *convert* the primary information to the intermediate, RNA. The division of eukaryotic genes into exons and introns requires subsequent *processing* of the primary transcript by the spliceosome^42 43^ before *export* of the RNA to the cytosol where it is *translated* to protein. The next step in cellular information flow is one of logistics, *deploying* the product to the right location and in the correct format. Vesicular transport in the cell, associated with the ER-Golgi apparatus, transports proteins to the cell surface and for secretion^15^, destinations that will be important as multicellular organisms gain in complexity and require signaling between cell types. Finally, the specific and regulated addition and removal of ubiquitin^44^, coupled with the proteasome, ensures that specific proteins are turned over, to provide a *limitation* to the pathway of information flow^45 46^.

We have also identified a number of genes whose function can be ascribed to maintaining the integrity of the information flow. These include DNA repair functions that are important for the stored information and processes such as ERAD that take place in the endoplasmic reticulum to remove defective proteins^27^. Unsurprisingly, information flow needs to be coordinated to the cell cycle and a number of genes that function in both have been identified (Fig. 6 and Supplementary data 7).

We can speculate that the increased range of functions of key proteins at each stage increases the efficiency of information flow through processes of neo- and sub-functionalization of paralogues and isoforms^47^. Whilst this paper identifies a general trend, further analysis of specific genes will be required to determine precisely how these efficiencies are achieved. It is clear, though, that the genes identified in this analysis do not greatly affect the cell-type specific activities of different differentiated cell types, rather they relate to the processing of information to generate those specific activities. Applying the concept of information flow within the cell is important because it will provide the context for a more detailed analysis and understanding of animal complexity.

## Acknowledgements

We would like to thank members of the Science Faculty, University of Portsmouth, for critical comments. JD, DLC and CS are grateful for University of Portsmouth Master’s support, a PhD bursary with Professor A. Callaghan and research support, respectively.

## Contributions

JD collected and analysed the data, DLC analysed the data and CS analysed the data and wrote the manuscript.

## Data availability

The authors declare that all other data supporting the findings of this study are available in the paper, in its supplementary information files (S1 and S3-7 are attached at the end of this document) and S2 is available in the University of Portsmouth repository, PURE, at DOI: 10.17029/3d3b91c7-3ae2-4ef8-9703-852709d04a8f and (S2-7) at https://github.com/colin17/complexity.git.

## Code availability

The code used, in R, to extract data from the Ensembl database is freely available at https://github.com/JackDean1/OrganismComplexity.

## Supplementary information

**S1.**
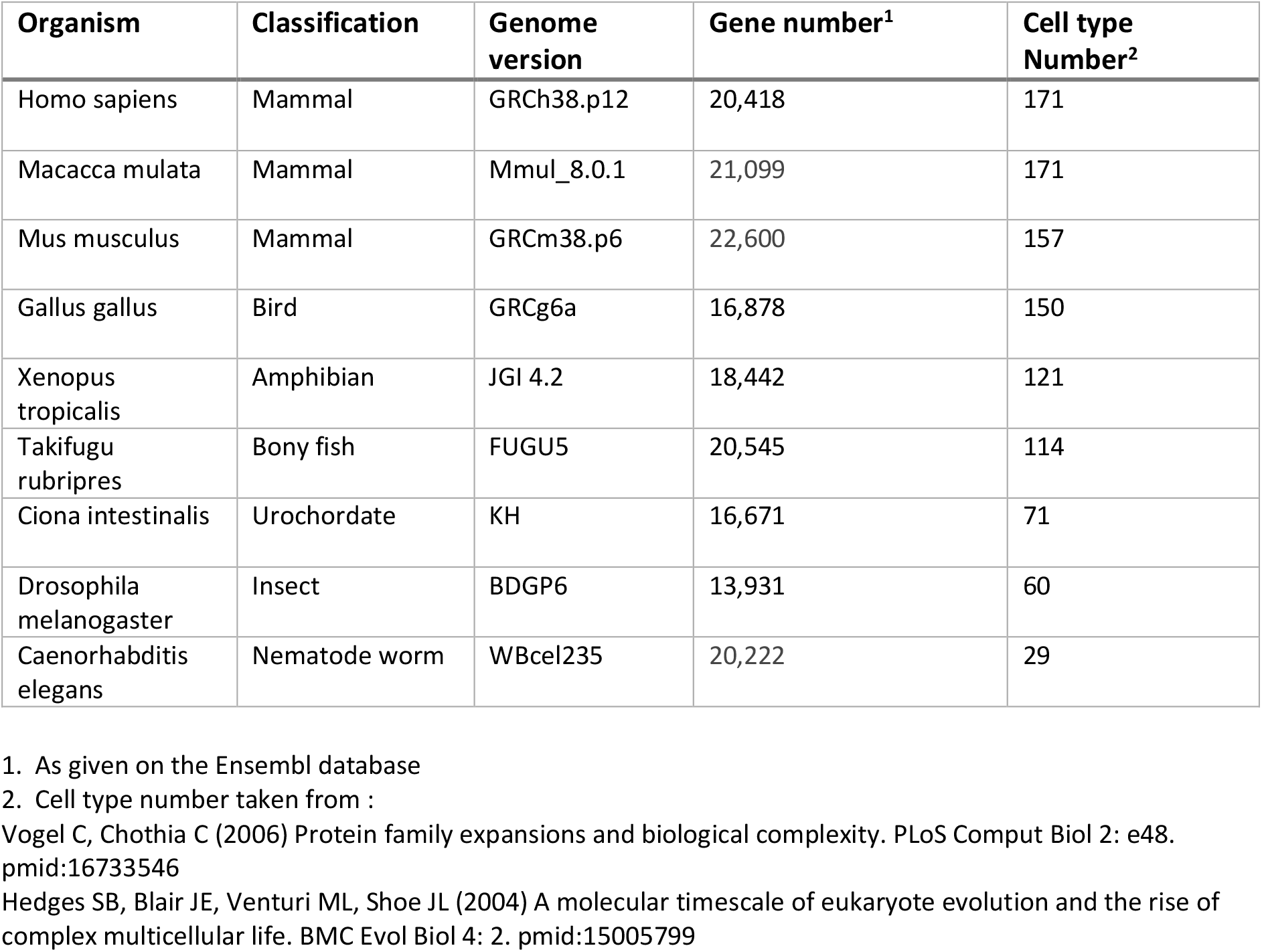
Table of the genome data used in this research.

**S2** Table of calculations of functional diversity for all genes across the eight species (CSV format).

**S3.**
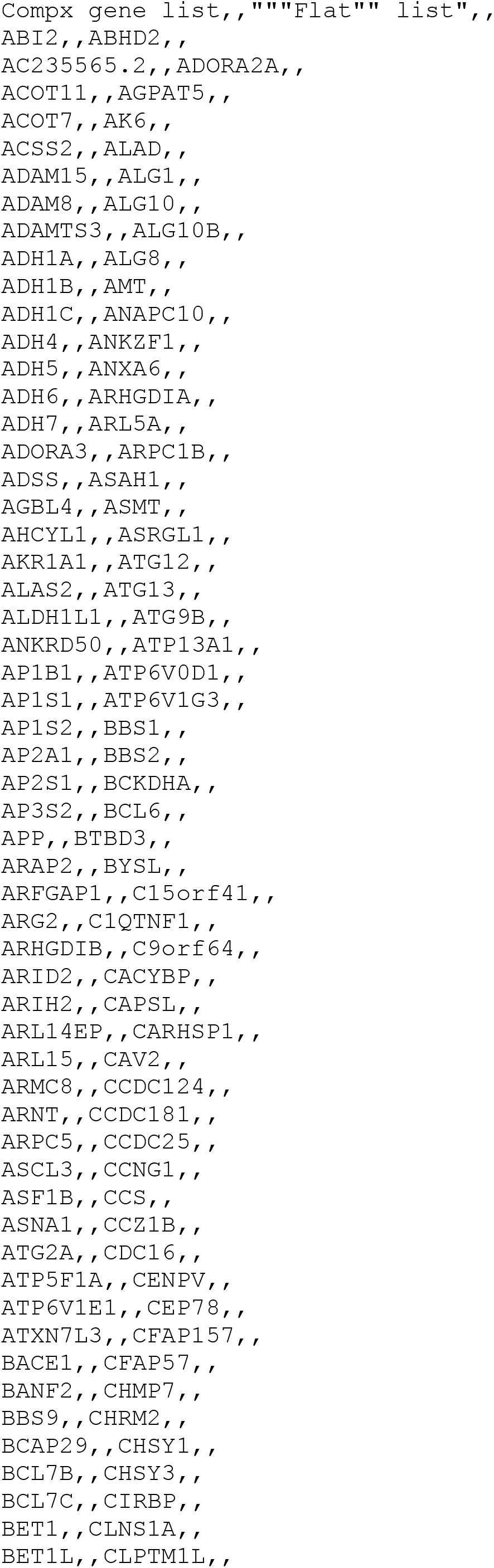
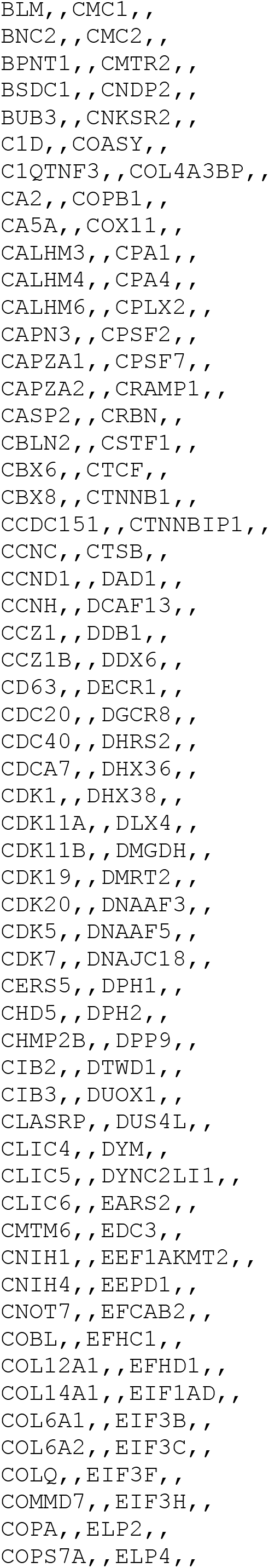
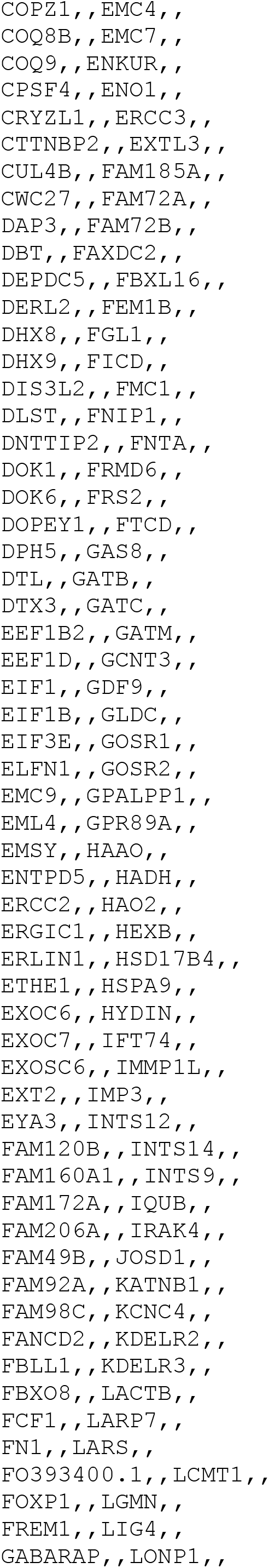
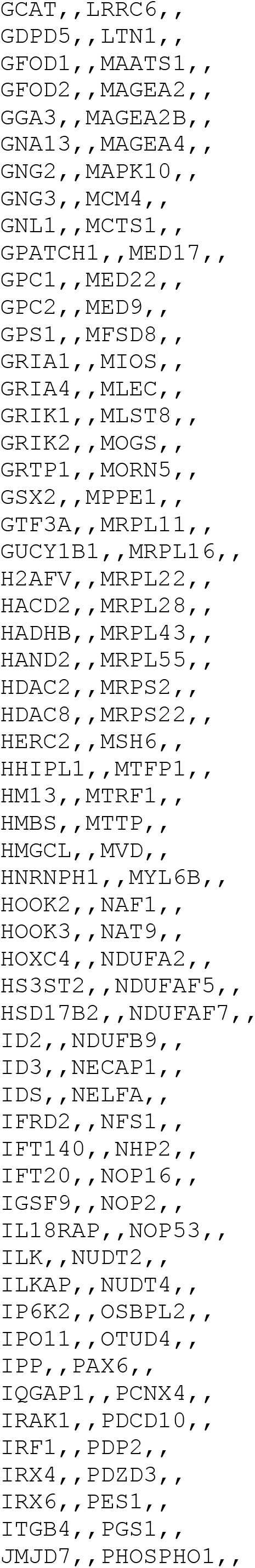
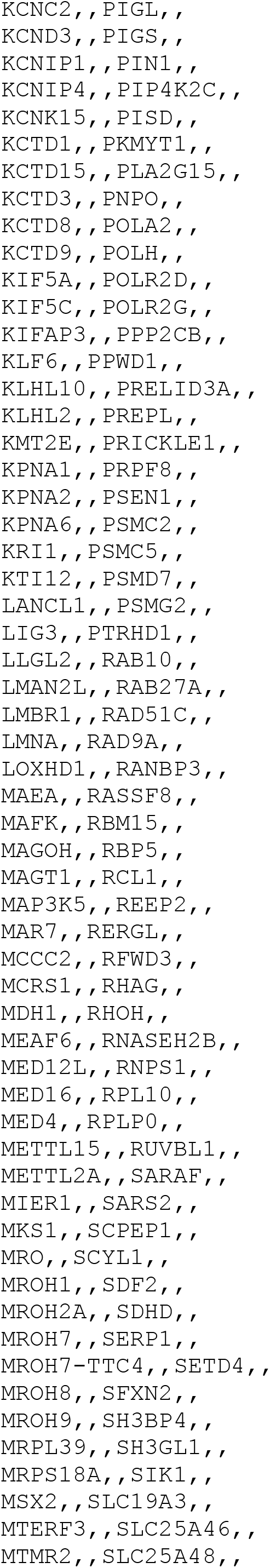
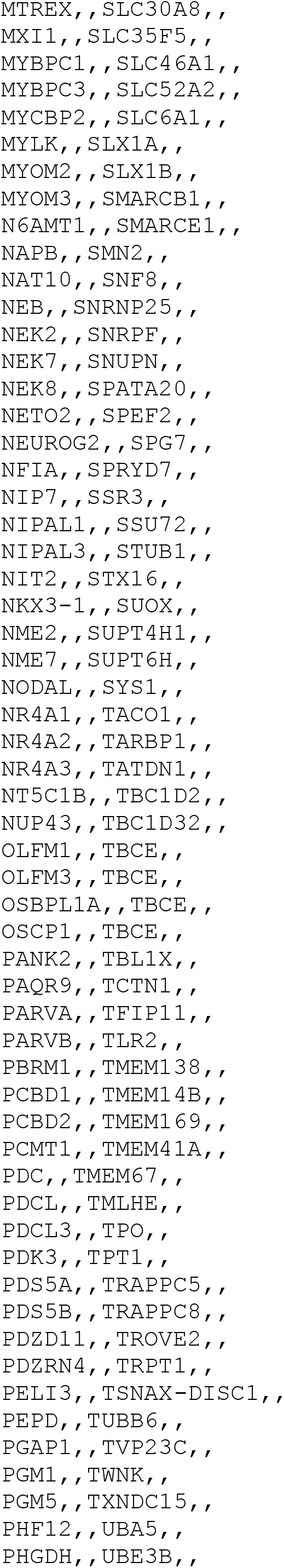
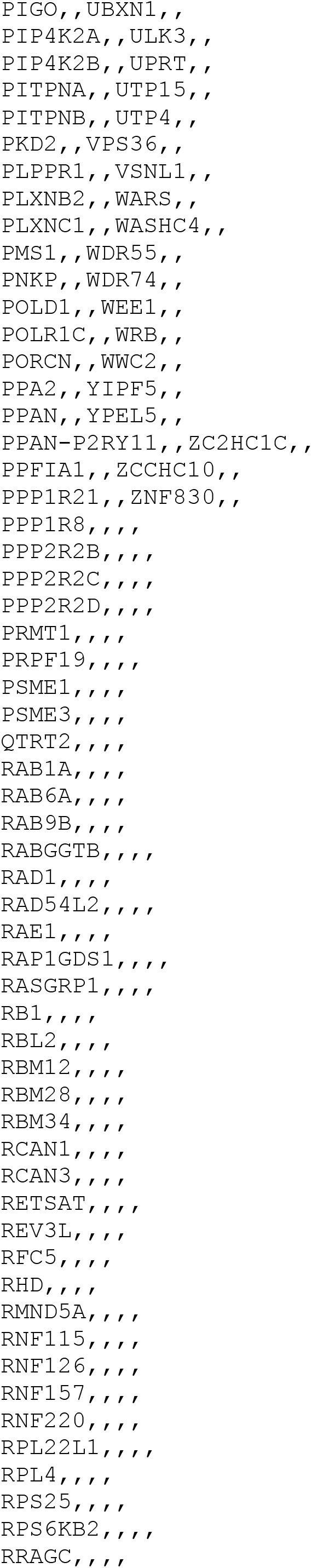
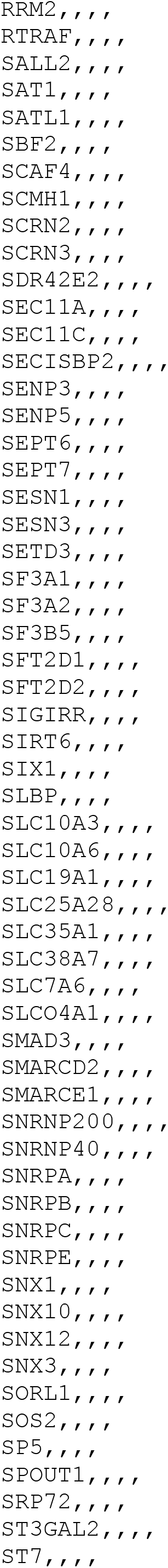
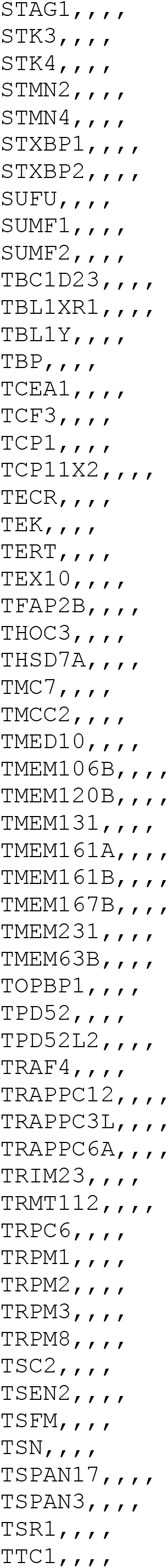
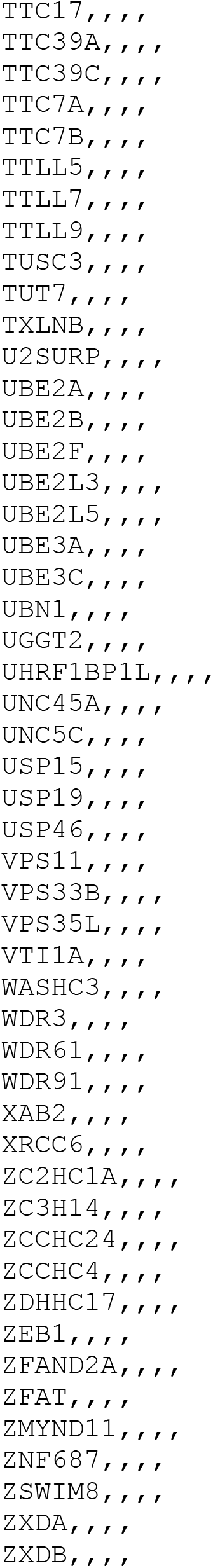
The Compx list of genes (CSV format)

**S4.**
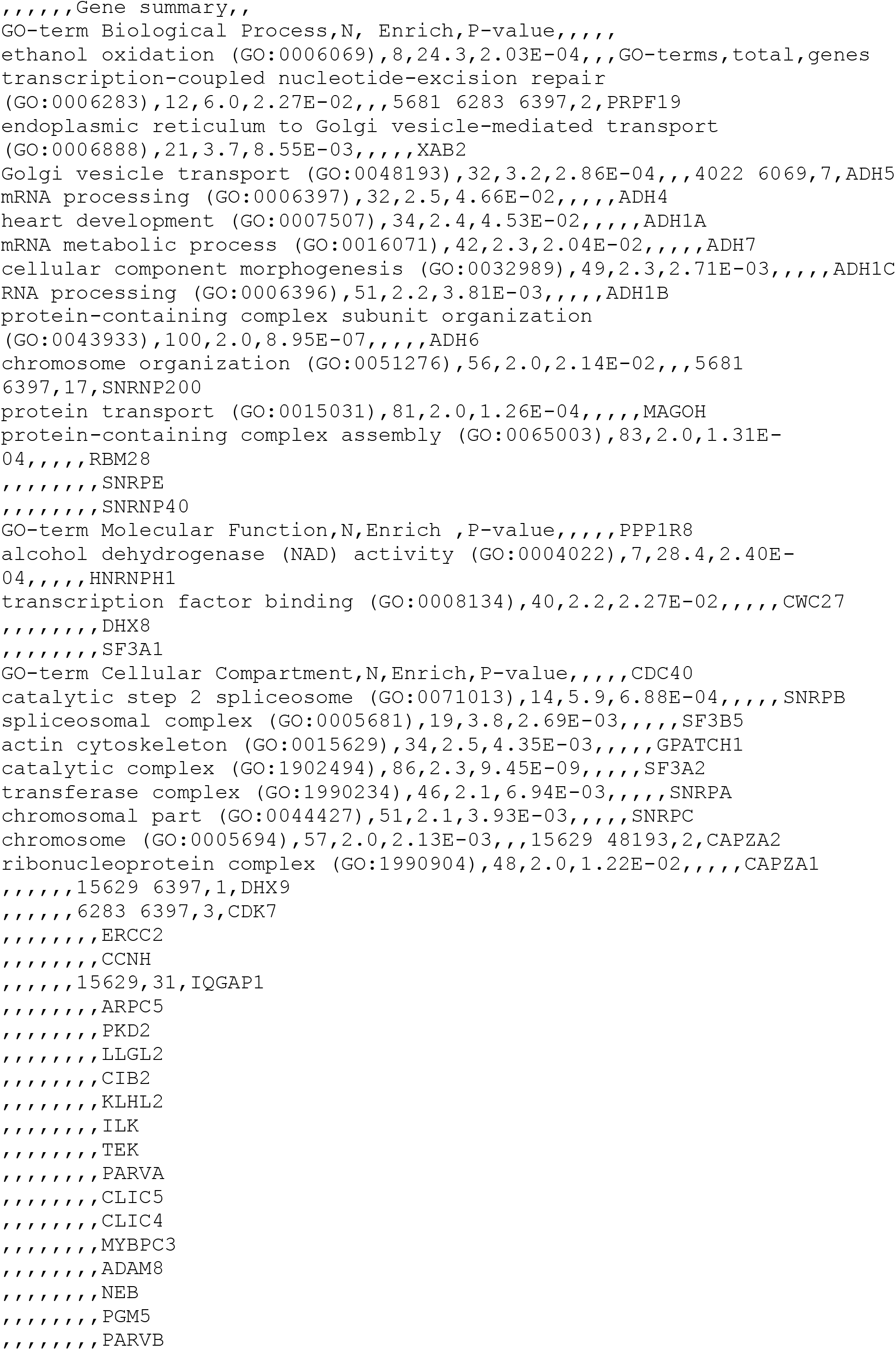
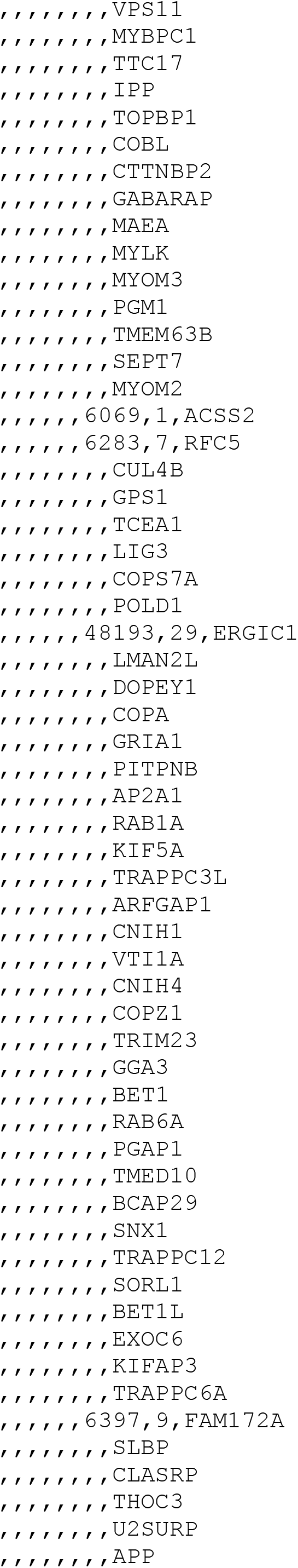
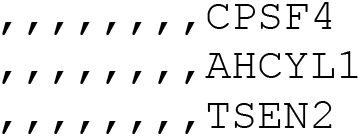
GO-term details for the Compx gene list (CSV format)

**S5.**
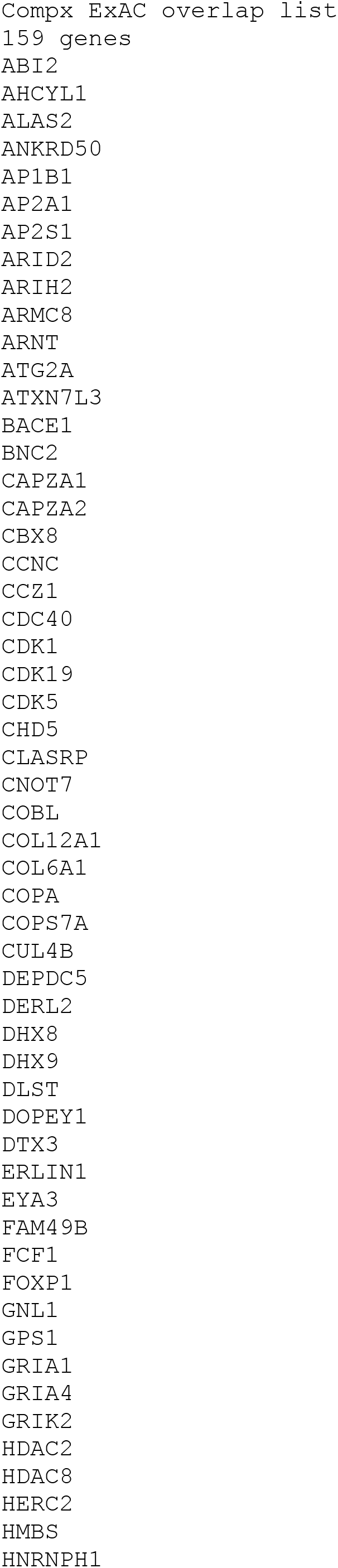
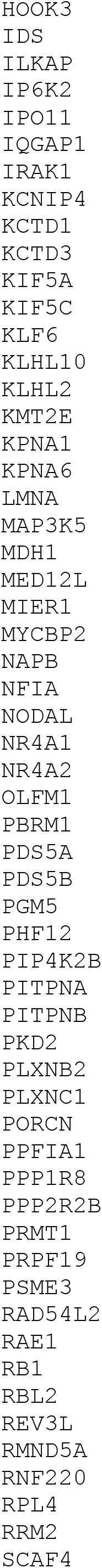
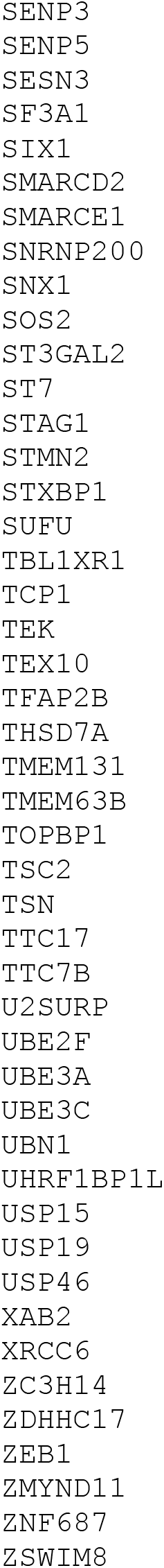
The Compx-ExAC overlap list of genes (CSV format)

**S6.**
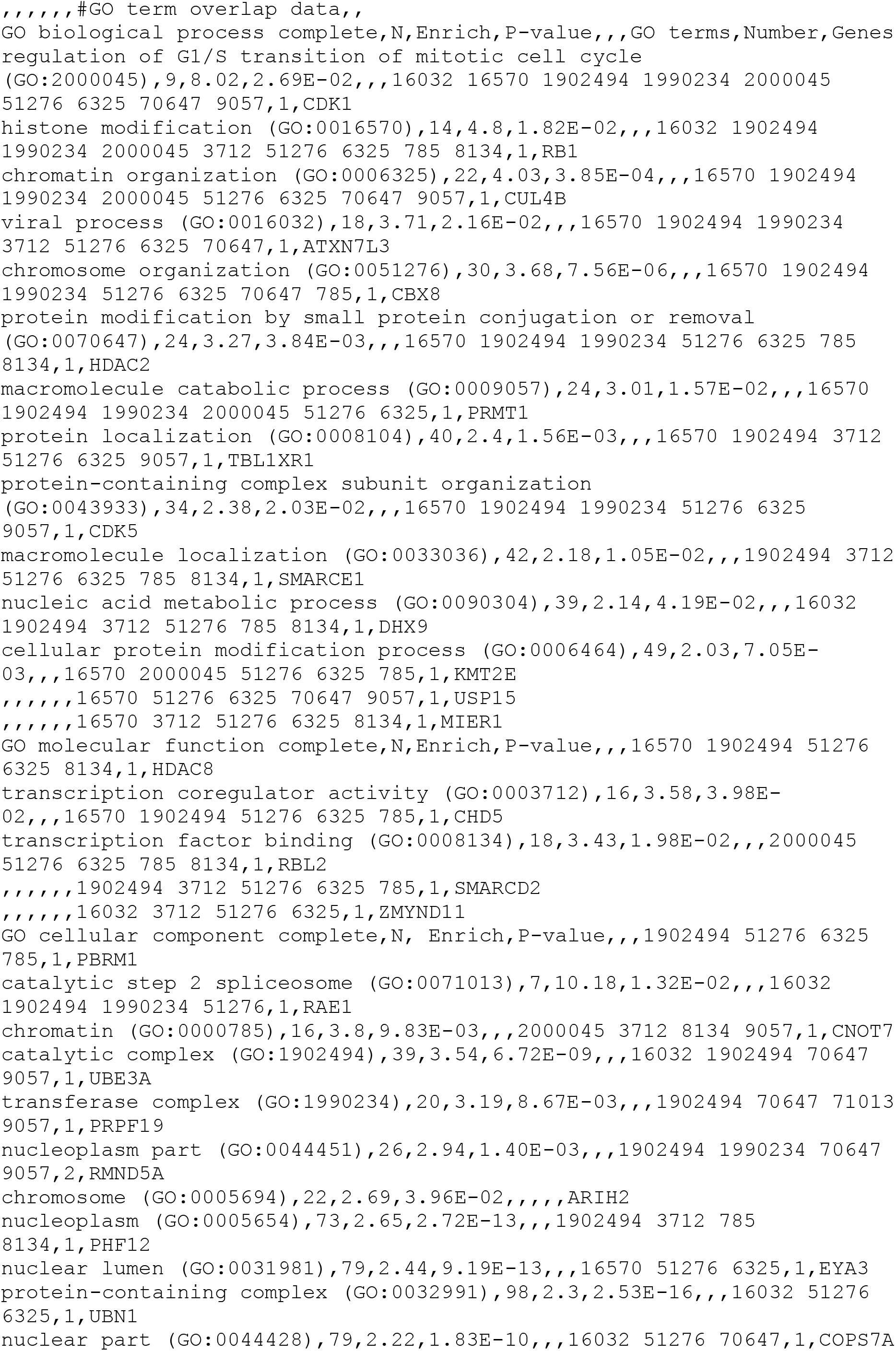
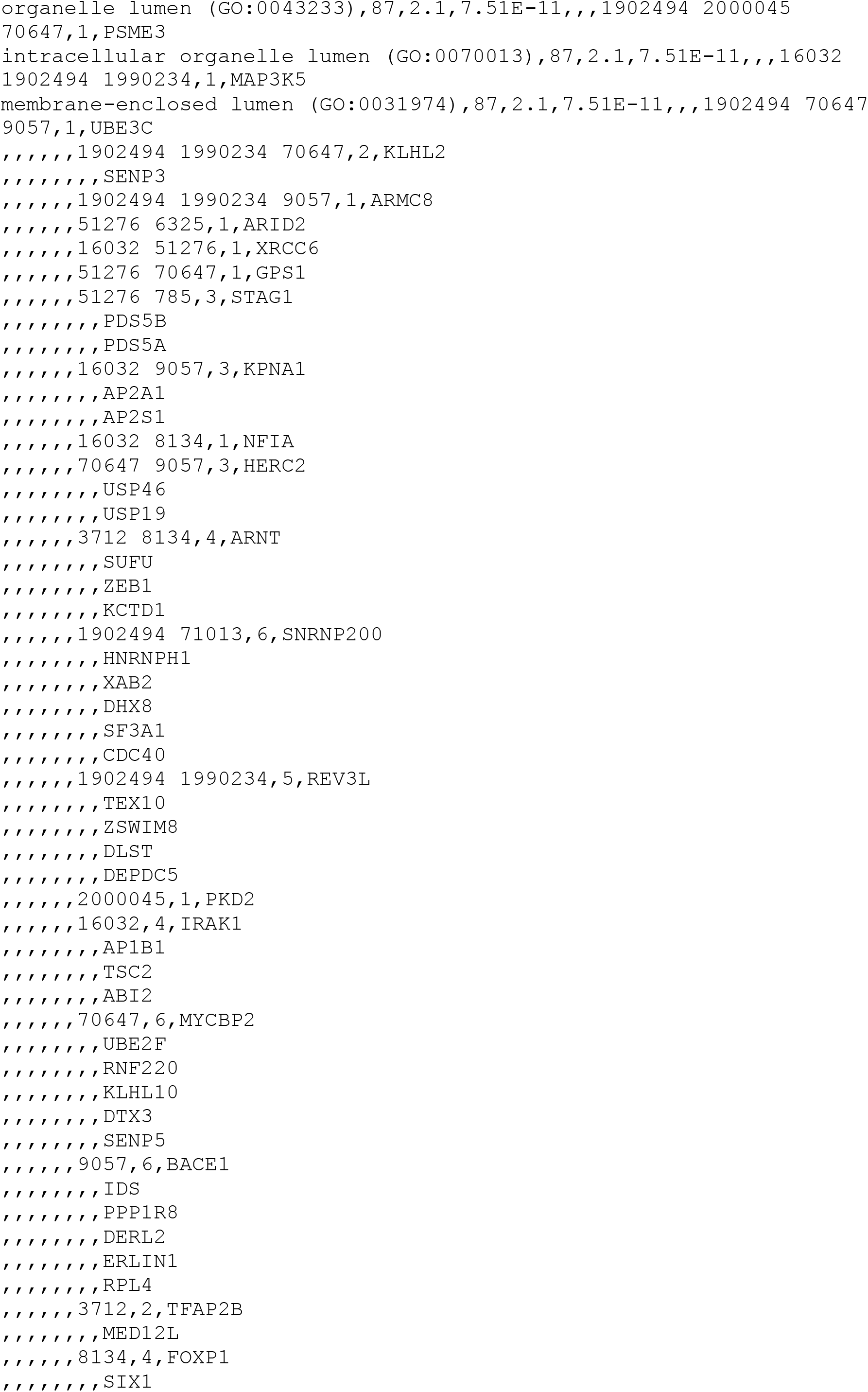
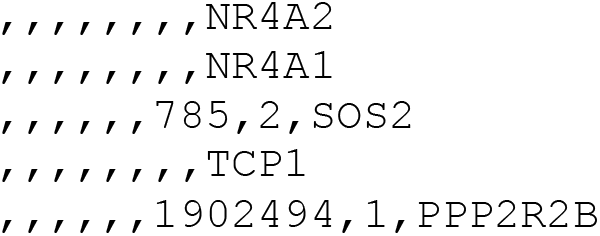
GO-term details for the Compx-ExAC overlap gene list (CSV format)

**S7.**
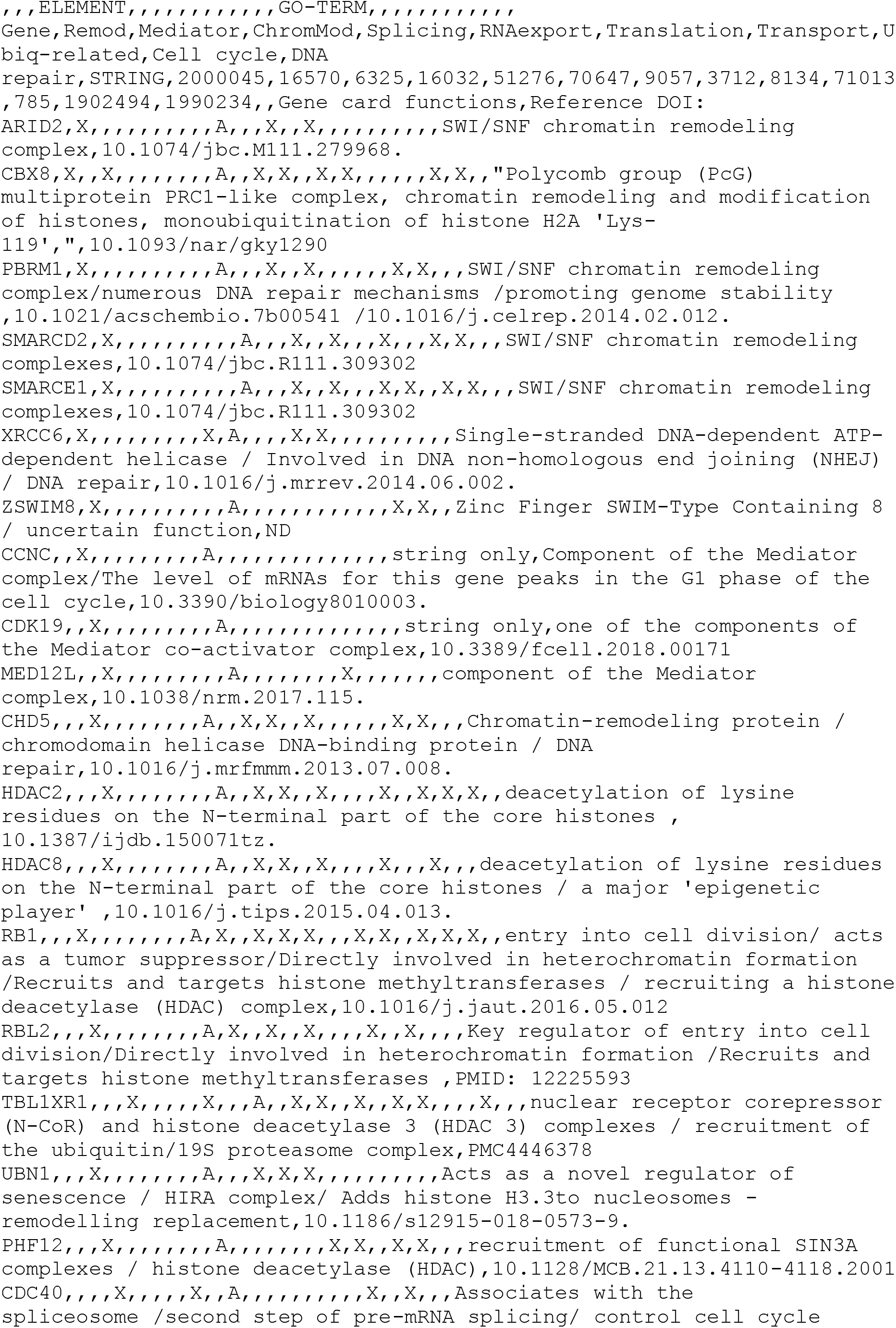
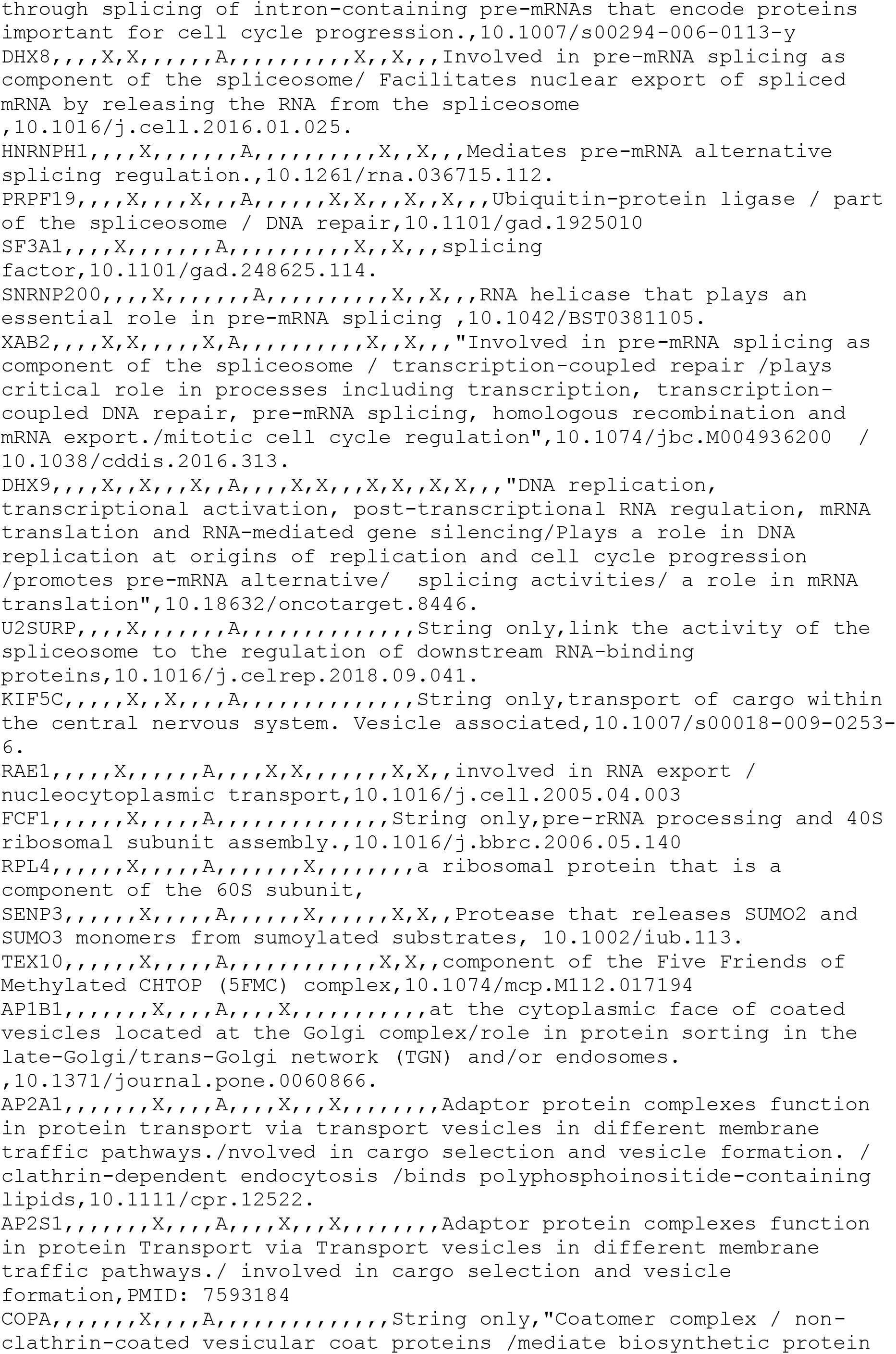
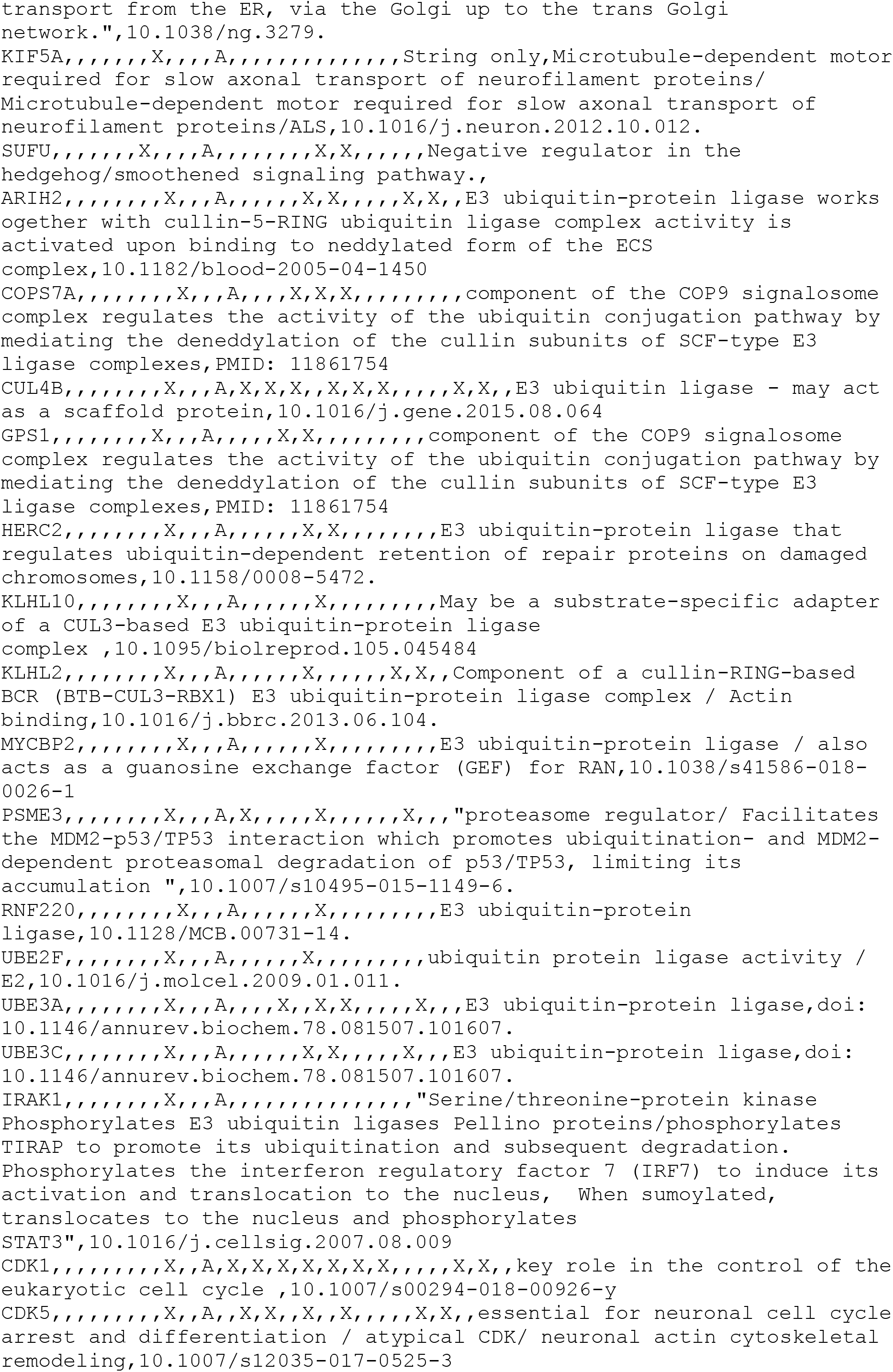
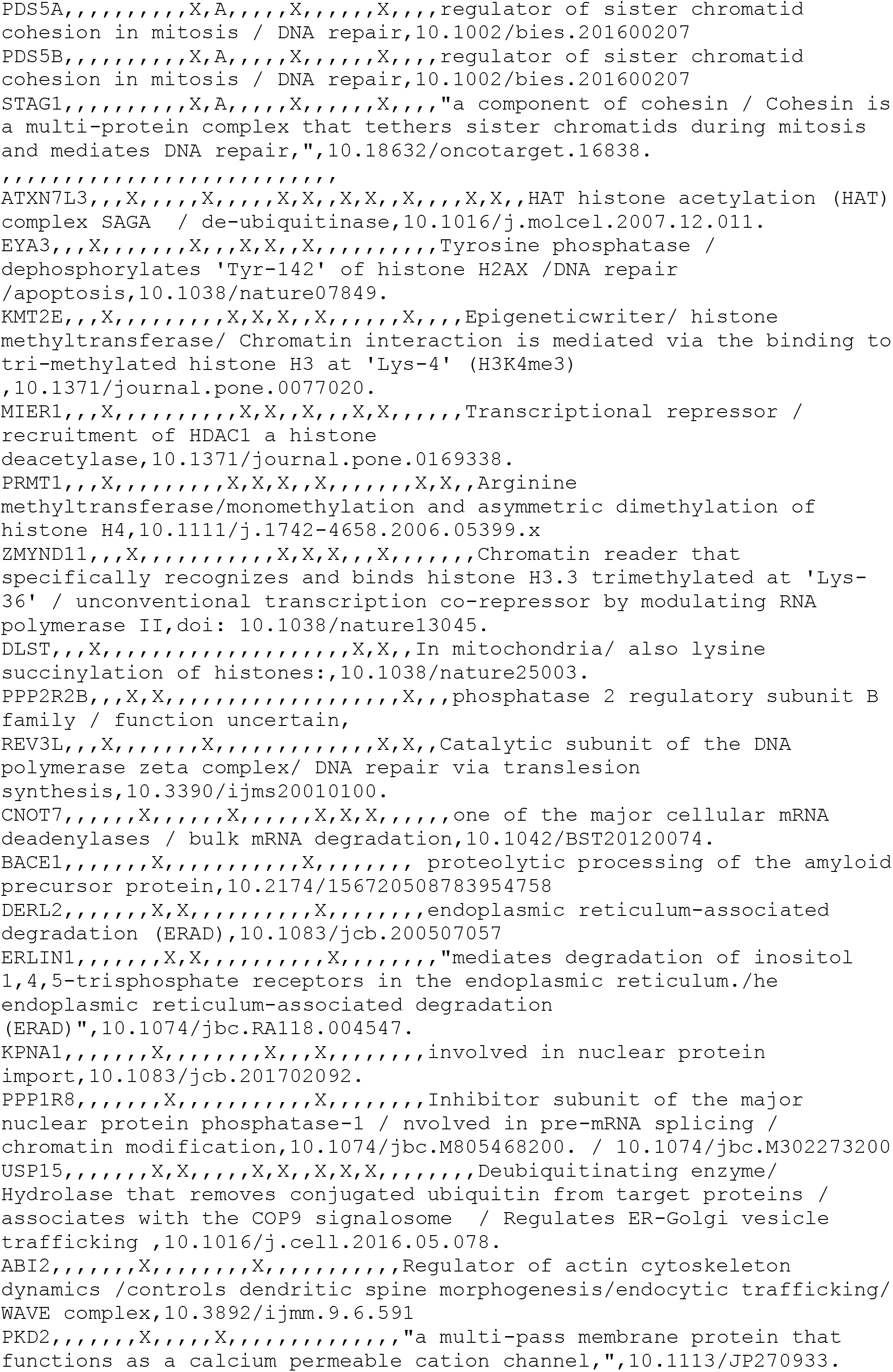
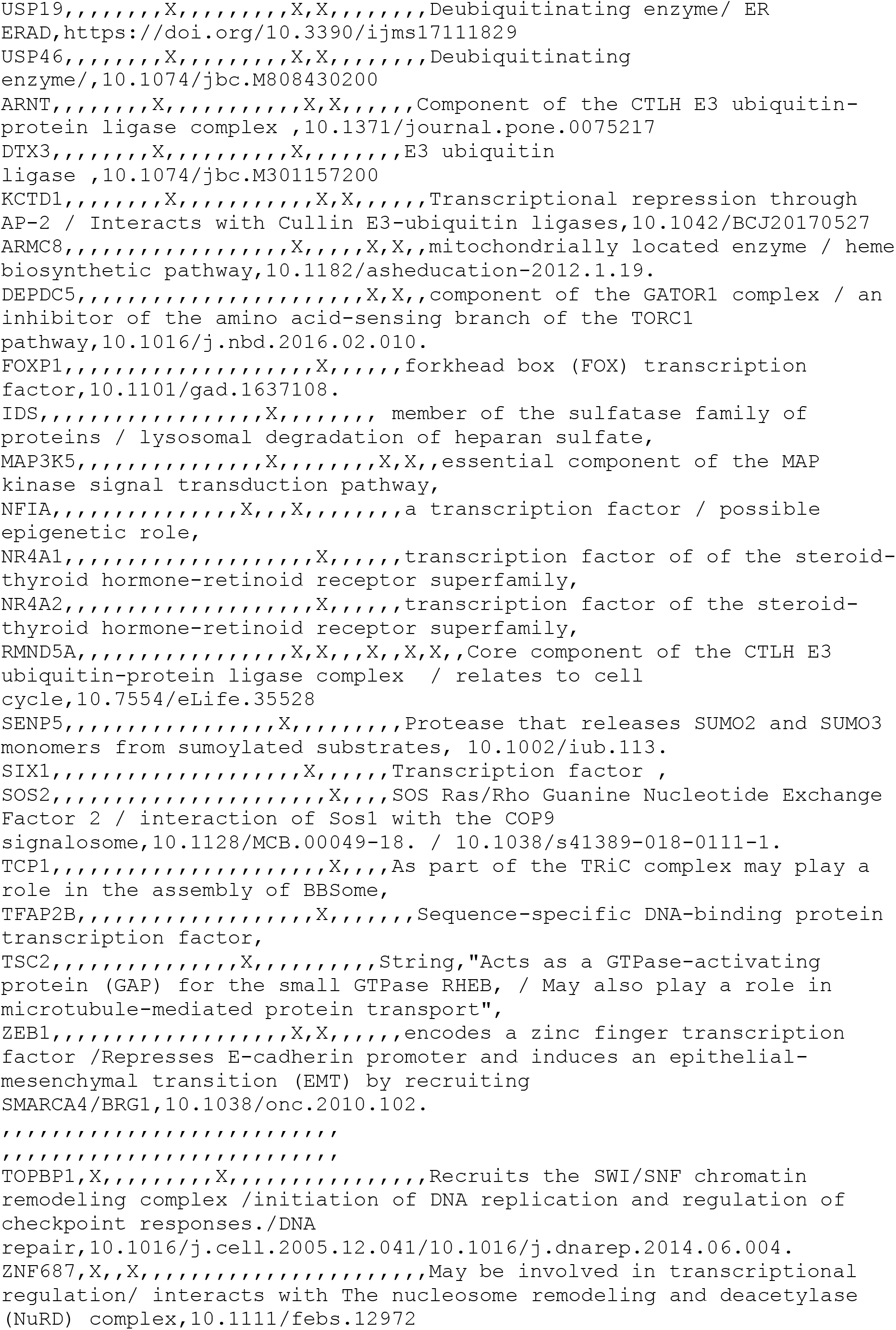
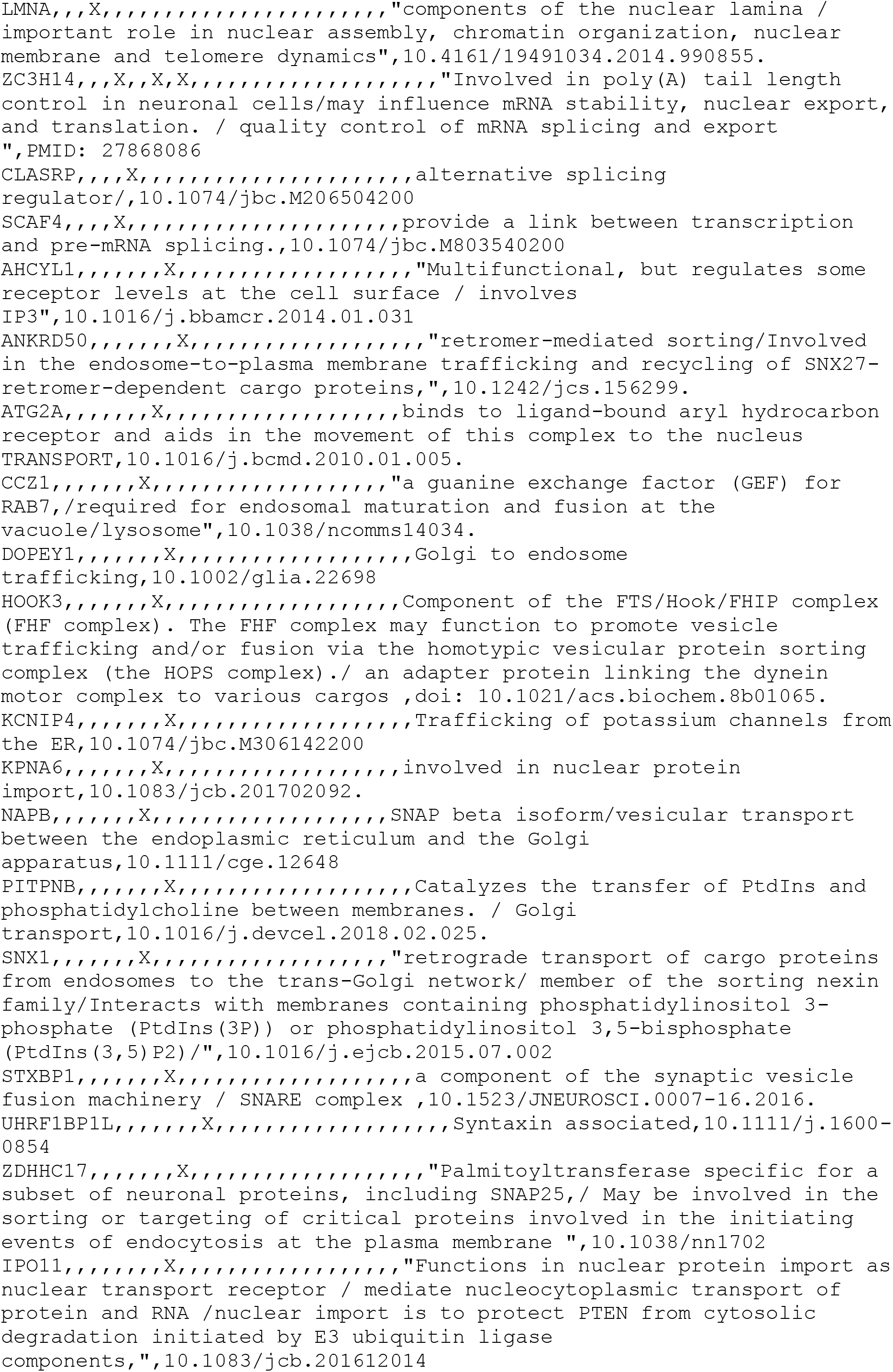
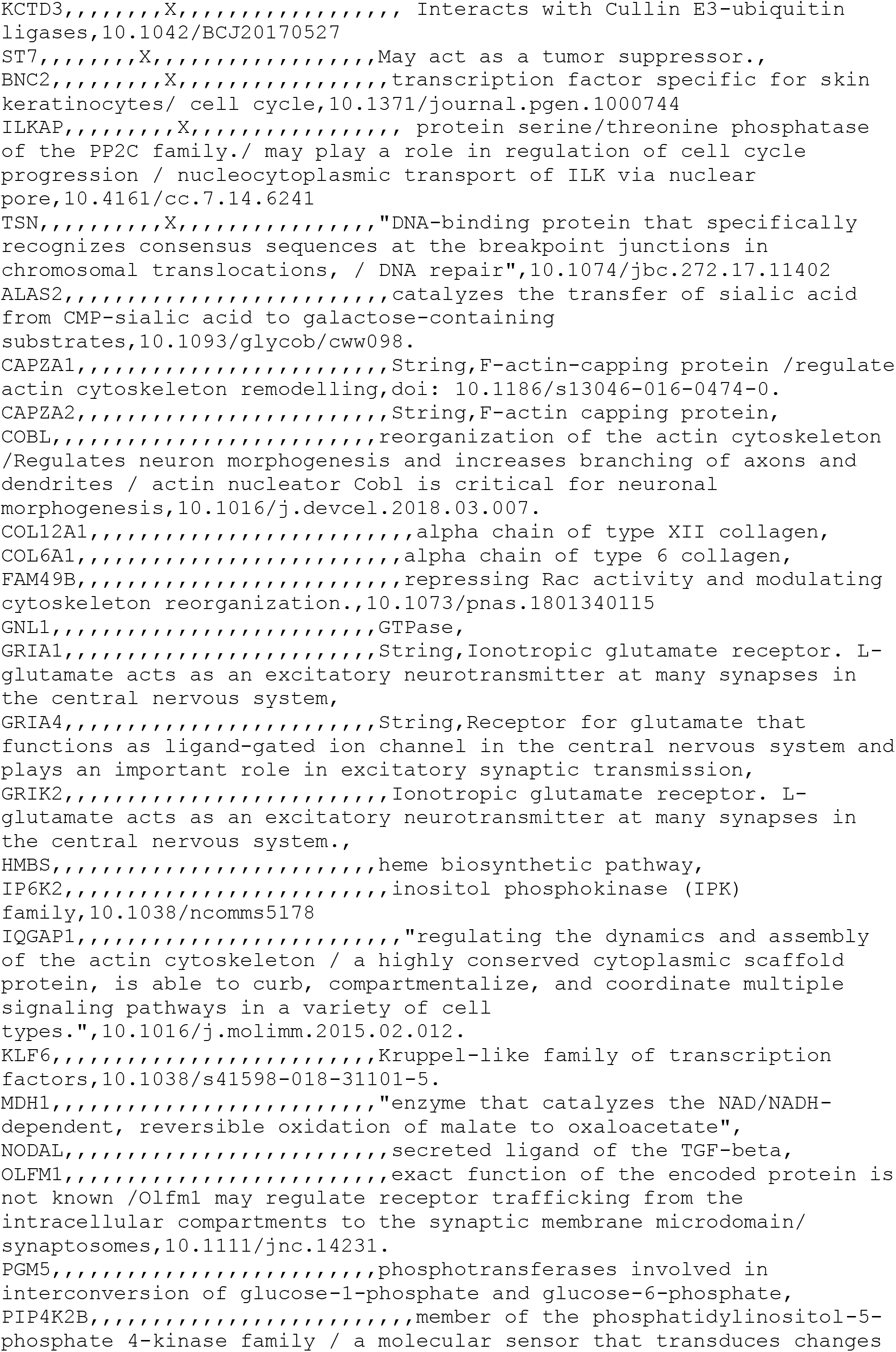
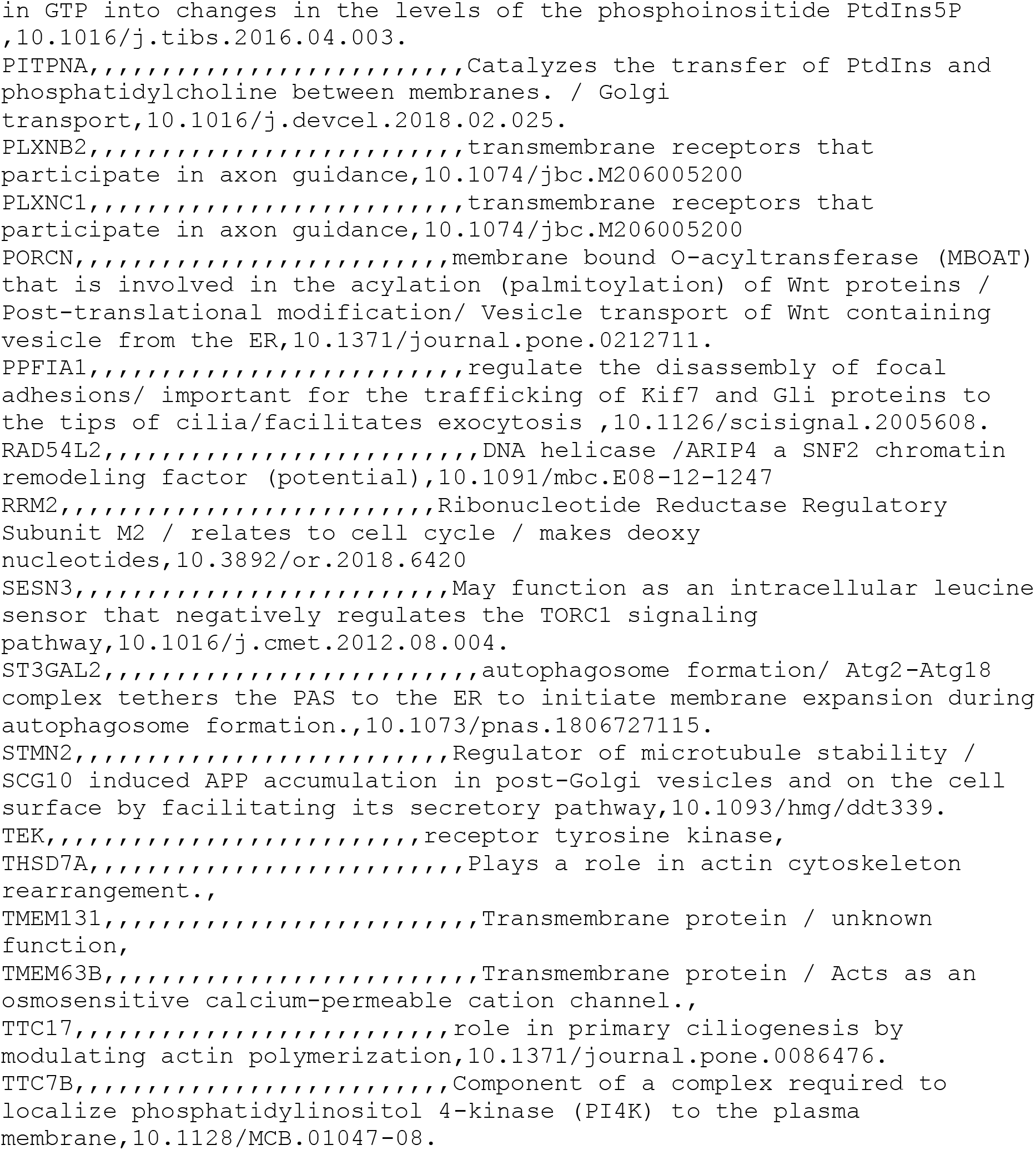
Compx-ExAC overlap gene list, functional details (CSV format)

## Supplementary data

Supplementary Data S1, S3-7

